# Glial Reactivity and Cognitive Decline Follow Chronic Heterochromatin Loss in Neurons

**DOI:** 10.1101/2022.08.29.505641

**Authors:** A.G. Newman, J. Sharif, P. Bessa, S. Zaqout, J. Brown, D. Richter, R. Dannenberg, M. Nakayama, S. Mueller, T. Schaub, S. Manickaraj, P. Böhm-Sturm, O. Ohara, H. Koseki, P.B. Singh, V. Tarabykin

## Abstract

In aging cells and animal models of premature aging, heterochromatin loss coincides with transcriptional disruption including the activation of normally silenced endogenous retroviruses (ERVs). Here we show that loss of heterochromatin maintenance and de-repression of ERVs results in a chronic inflammatory environment characterized by neurodegeneration and cognitive decline. We discovered differential contributions of HP1 proteins to ERV silencing where HP1γ is necessary and sufficient for H4K20me3 deposition and HP1β deficiency causes aberrant DNA methylation. Combined loss of HP1β and HP1γ resulted in loss of DNA methylation at ERVK elements. Progressive ERV de-repression in HP1β/γ DKO mice was followed by stimulation of the integrated stress response, an increase of Complement 3+ reactive astrocytes and phagocytic microglia. This chronic inflammatory state coincided with age-dependent reductions in dendrite complexity and cognition. Our results demonstrate the importance of preventing loss of epigenetic maintenance, as this will be the only way postmitotic neuronal genomes can be protected and/or renewed.

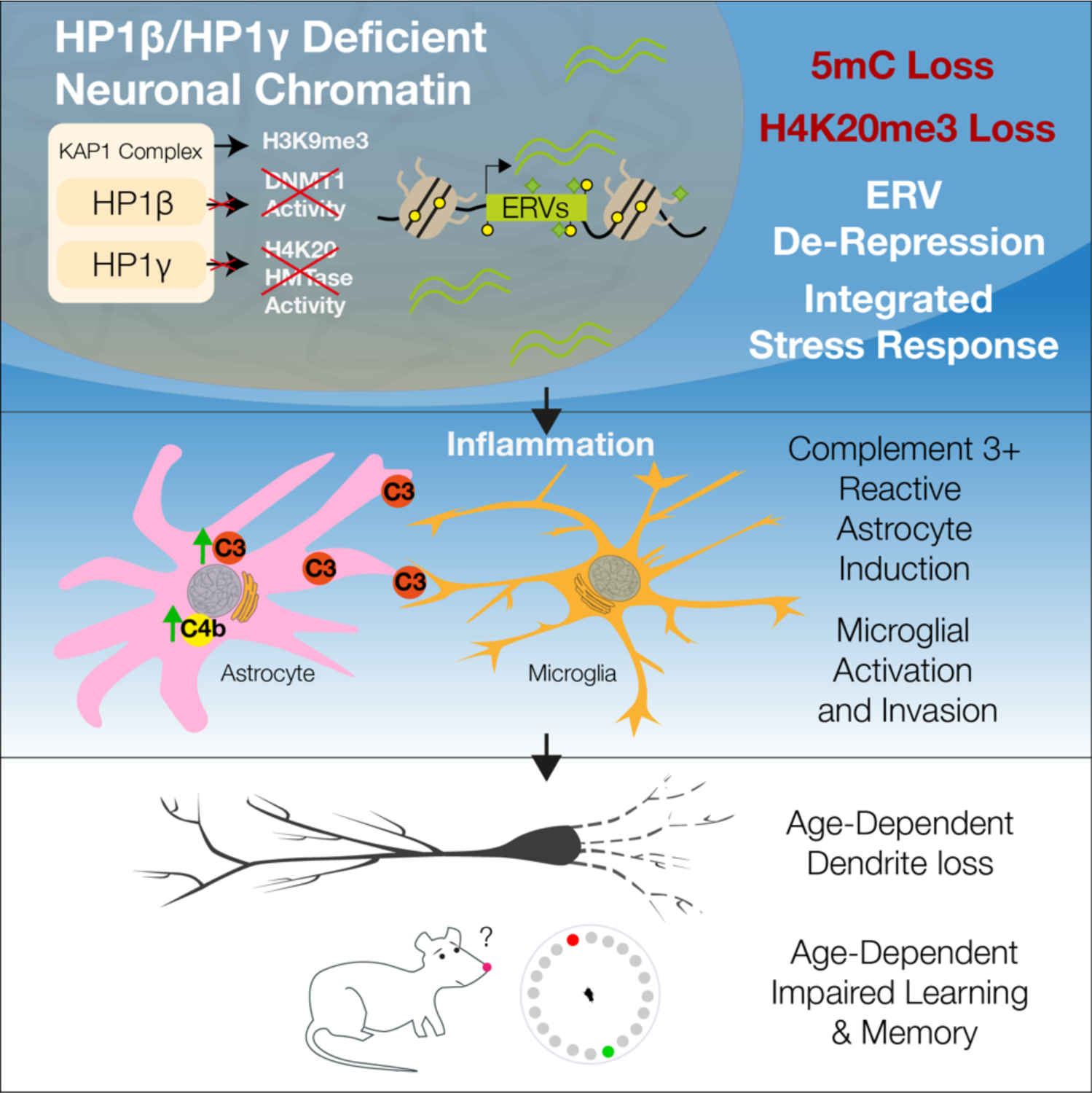

## Introduction

Aging neurons operate under conditions of high cellular stress without renewal by cell division. Under normal physiological conditions, neurons continuously break their own DNA at enhancer regions ^1,2^, and their high metabolic rate results in an excess of reactive oxygen species harmful to protein and DNA integrity^3^. Neuronal genes most sensitive to oxidative DNA damage are downregulated with age^4^, resulting in either cell death or cascading dysfunction and de-differentiation^5^.

Little is known about what occurs in heterochromatin in aging neurons. However, in other cell types and human progeria models, a loss of DNA methylation^6,7^, histone 3 lysine 9 tri-methylation (H3K9me3) and associated proteins is observed with age^8–10^, along with a decreased association of heterochromatin with the nuclear lamina^11^. These age-related changes result in the activation of normally silenced repetitive elements such as endogenous retroviruses (ERVs)^12^ and Long Interspersed Nuclear Elements (LINEs)^10^. Elevated transcription of ERVs have been observed in pathological states such as exogenous viral infections^13,14^, cancer^15^, neurodegeneration^16,17^, multiple sclerosis^18,19^, ALS^20^, and Alzheimer’s disease^17^. Elevated levels of ERVs have also been seen in models examining factors associated with neurodegenerative diseases such and Tau^21^ and TDP-43^22^, while α-synuclein has been shown to affect chromatin and the maintenance of ERVs directly^23,24^. However, a causal relationship between ERVs and the initiation of neurodegeneration has yet to be determined.

ERVs are silenced by the KAP1 repressor complex which recruits histone deacetylases, DNA methyltransferases, and histone methyltransferases and several cofactors to induce heterochromatin formation (reviewed in ^25^). Here, the histone methyltransferase SETDB1 catalyzes H3K9me3 methylation^26^, which serves as a high-affinity binding site for Heterochromatin Protein 1 (HP1)^27,28^, which facilitates compaction and silencing^29^.

We mimicked age-related heterochromatin loss by deletions of members of the HP1 family in the mouse brain. Unlike other mutants of enzymatic epigenetic modifiers—which typically have severe developmental phenotypes—the removal of HP1 proteins mimics the destabilization normally seen in aged cells, whose lower levels of H3K9me3 naturally result in less HP1 binding, activity, and stability. In doing so, we describe the molecular contributions of HP1β (*Cbx1*) and HP1γ (*Cbx3*) to ERV silencing and uncover an endogenous cause of the known age-related increases^30,31^ of Complement in the brain.

All three HP1 homologs are robustly expressed in post-mitotic neurons (fig S1a,b). To test whether heterochromatin loss can drive neuronal aging *in vivo*, we engineered mice to be conditionally deficient for HP1β and HP1γ in the cerebral cortex using the *Emx1^Cre^* deleter mouse strain. We observed a partial malformation of the infrapyramidal blade of the dentate gyrus (DG) due to depletion of Ki67+ progenitors following loss of HP1β (fig **S1c,d**), which is consistent with HP1β’s established role in mitotic stability^32^. Apart from this minor developmental defect, we detected no overt changes in cortical cytoarchitecture in HP1 single and double mutants (fig. **S1e,f**), making these mutants a suitable model to study the long-term effects of HP1-related heterochromatin loss.

## Results

### HP1 deficiency results in de-repression of ERVs and an innate immune response

Given the known role of HP1 proteins in maintaining repression of non-coding elements ^33^, we performed cRNA-RNA *in situ* hybridization for the murine endogenous retrovirus (ERV) Intracisternal Alpha Particle (IAP) and could observe its robust de-repression in the HP1β/γ DKO, which was especially strong in the hippocampus and absent from the dentate gyrus (fig. **1** & **S2b**). RNAseq on young and aged hippocampi (fig. 1a) confirmed that repeats and chimeric transcripts were de-repressed in HP1β/γ DKO (fig. **1b****,c**, **S2c,d** & Data S1). The same analysis revealed that apart from repeats and chimeric transcripts, the primary sources of genotype-dependent variance fell into three further categories (fig. **1a**, Data S1): The first are dentate gyrus related genes underrepresented in HP1βKO and HP1β/γ DKO (140) such as Prox1, Dsp, Trpc6, Plk5 and Cdh9 (Data S1). The remaining two categories were upregulated (549): first, expression of the entire protocadherin cluster (cPcdh) was elevated in HP1γKO and HP1β/γ DKO (which is further observed in gene ontology analyses (fig. **S2e**)), as were a small subset of canonical and non-canonical imprinted genes (Data S1). Second, in HP1β/γ DKO we observed changes in genes related to inflammation including innate immune pathways and unfolded protein response (UPR) (fig. **1b**).

**Figure 1.**
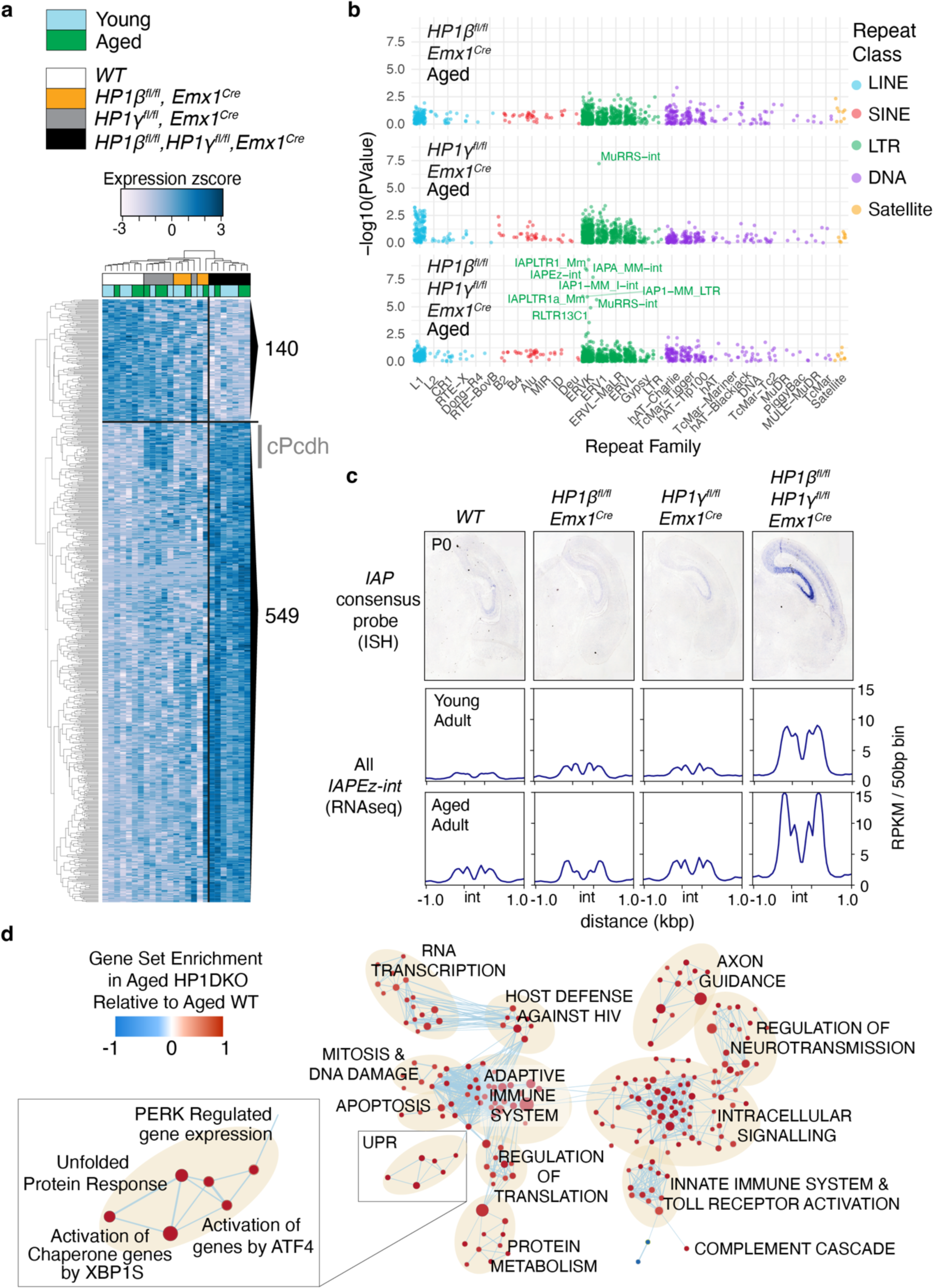
Loss of both HP1β and HP1γ results in activation of noncoding elements and induction of the integrated stress response. **a** Genes (inc. repetitive elements) significantly changed in young and aged *HP1β^fl/fl^HP1γ^fl/fl^Emx1^Cre^* hippocampi (689 genes, corrected p <0.05). **b** Pseudo-Manhattan plot of significant changes to repetitive element transcription in aged HP1 mutants. Repeats from the ERVK subfamily, which contains the evolutionarily recent *IAP*s, is most strongly affected in *HP1β^fl/fl^HP1γ^fl/fl^Emx1^Cre^* hippocampi. **c** *In situ hybridization* using a consensus probe for IAP in P0 brains and RNAseq read coverage over the IAPEz internal fragment in young and aged adults (read coverage is RPKM normalized reads per 50bp bin). Gene Set Enrichment of upregulated transcripts **d** shows several affected pathways including host defense, Toll Receptor activation, unfolded protein response and the complement cascade.

### HP1γ is required for H4K20me3 deposition and regulation of Protocadherin genes

To understand the epigenetic pathway by which HP1β and HP1γ regulate repression of ERVs, we investigated the effect of HP1β and HP1γ mutations on the HP1-related histone modifications H3K9me3 (histone 3 lysine 9 trimethylation) and H4K20me3 (histone 4 lysine 20 trimethylation). H3K9me3-bound HP1 can recruit Suv420h1/2 HMTases and direct local H4K20me3 deposition^34^, and both histone modifications are abundant in wt post-mitotic neurons (fig. **2a****, S3a**). We found that H3K9me3 remained unchanged (fig. **2a**) but observed a specific loss of H4K20me3 in specifically HP1γ-deficient neurons(fig. **2a****, S3b**), consistent with previous observations in spermatocytes^35^. We also found that HP1γ was sufficient for H4K20me3 deposition because re-addition of HP1γ to HP1β/γ DKO cortices by *in utero* electroporation at E14 restored H4K20me3, albeit not to the levels seen in adjacent interneurons where HP1γ is not deleted (fig. **2b**). The H3K9me3-HP1γ-H4K20me3 pathway also appears to regulate isoform selection at the protocadherin (cPcdh) cluster. While all protocadherin isoforms can be observed in bulk RNAseq, single neurons express a unique combination of protocadherin isoforms which is clonally defined during neurogenesis^36–38^. Thus, bulk ChIPseq shows the protocadherin cluster is marked with H3K9me3 and H4K20me3 (fig. **S3c**) corresponding to single cell silencing of unused exons observed at the population level. In the HP1γKO, H4K20me3 is lost and there is elevated expression of cPcdh genes (fig. **1a**, **S3c**). Given the known requirement of the HP1 chromoshadow domain (CSD) for association with the H4K20me3 HMTase Suv420h2^39^, we carried out co-immunoprecipitation and co-localization analysis to identify residues in the CSD essential for the interaction of HP1γ with Suv420h2 (fig. **2c-e**, S4). Given that the cPcdh cluster is regulated by the SETDB1 HMTase that generates H3K9me3^40^, it seems likely that a H3K9me3-HP1γ-H4K20me3 pathway might be an important mechanism for regulating chromosomal domains, such as the protocadherin cluster and clustered retrotransposons.

**Figure 2.**
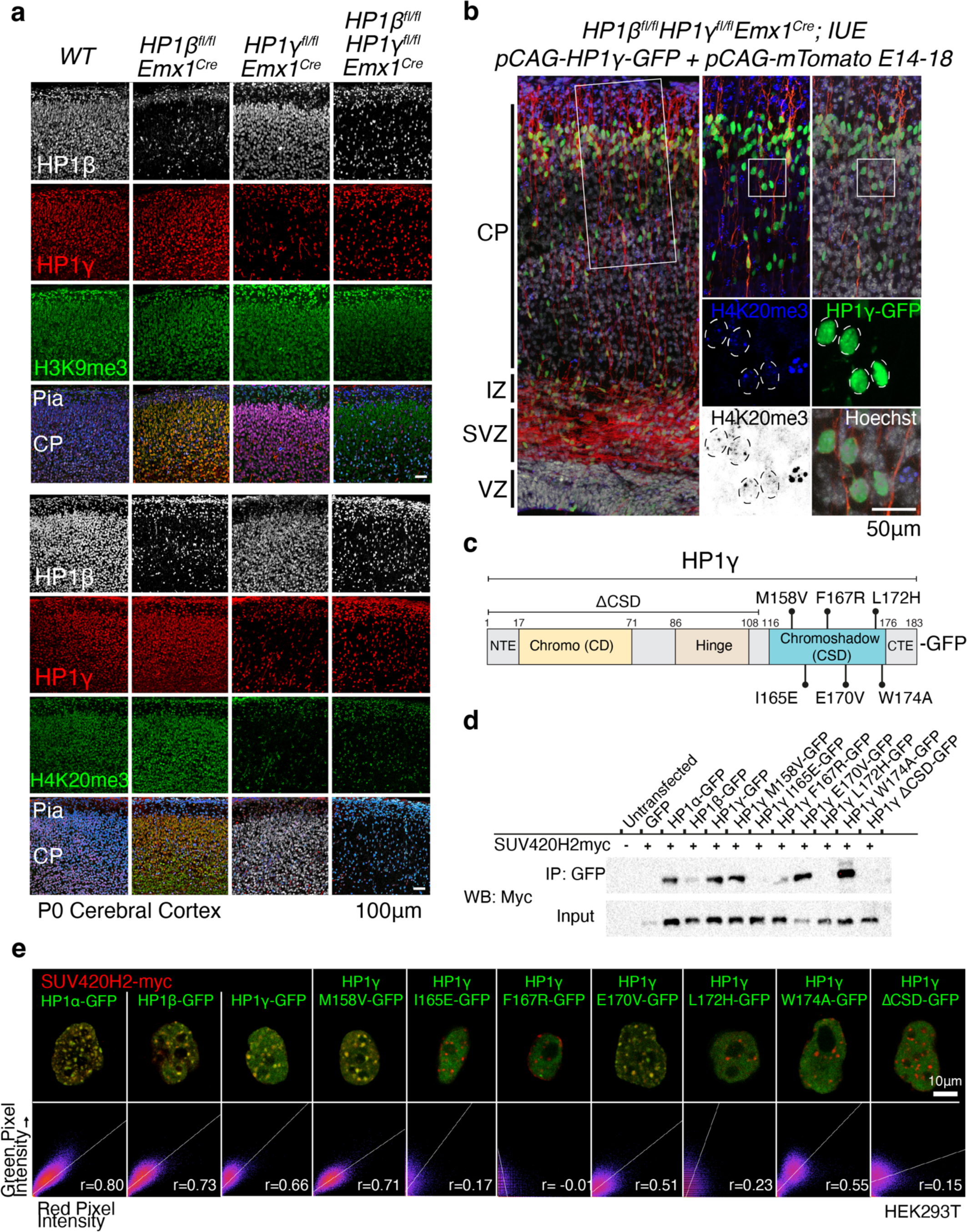
HP1γ is necessary and sufficient for deposition of H4K20me3. **a** H3K9me3 levels are unchanged in HP1β and HP1γ deficient neurons. H4K20me3, is unaffected in wild-type interneurons but lost completely in HP1γ-deficient pyramidal neurons. **b** Re-addition of HP1γ into developing mutant brains by *in utero electroporation* can restore H4K20me3 (magnification, nuclei outlined by dashed circles). (CP = cortical plate IZ = intermediate zone, SVZ = subventricular zone, VZ = ventricular zone). **c** Schematic of HP1γ protein product with domains and point mutations tested. **d** Co-immunoprecipitation of SUV420H2 with HP1 proteins and point mutants confirms residues in the chromoshadow domain (CSD) of HP1γ are essential for its binding with SUV420H2. **e** Co-localization of C terminal GFP-tagged HP1α, HP1β, HP1γ and HP1γ mutants with C terminal myc-tagged SUV420H2.

### HP1β deficiency results in aberrant DNA methylation

Prominent increases in the expression of tissue specific imprinted genes (Data S1) in HP1βKO and HP1β/γ DKO mutants indicated that DNA methylation may be affected by HP1β deficiency.

We engineered an ES cell line that contains an ERT2-Cre transgene where all three HP1 genes are floxed. This system allowed for the deletion of all three HP1 genes following the addition of tamoxifen, (thus termed HP1cTKO) that we then be profiled for DNA methylation using reduced-read bisulfite sequencing (RRBS) and changes to KAP1, H3K9me3 and H4K20me3 (Figure **3a**). We found that triple deficiency of HP1 proteins results in reduced KAP1 at imprinting control regions (ICRs) marked by ZFP57 (fig. **S5a**), while DNA methylation at these ICRs including *Nnat* shows mixed changes (fig. **S5b,c**). KAP1 was unchanged over IAP elements in HP1cTKO ES cells, with a reduction in H3K9me3 and an expected absence of H4K20me3 (fig. **3b**). While ES cells are not expected to utilize protocadherins the same way as neurons, regulatory H3K9me3 in the cPcdh is unaffected in HP1cTKO ESCs despite H3K9me3 and H4K20me3 being lost at adjacent IAP elements (fig **S5d**, arrows). This additionally suggests that regulatory H3K9me3 is unaffected in *HP1γ^fl/fl^ Emx1^Cre^* mutants, and that it is disruption of the HP1γ-H4K20me3 pathway that causes elevated Pcdh expression in *HP1γ^fl/fl^ Emx1^Cre^* and subsequently *HP1β^fl/fl^ HP1γ^fl/fl^ Emx1^Cre^* brains.

**Figure 3.**
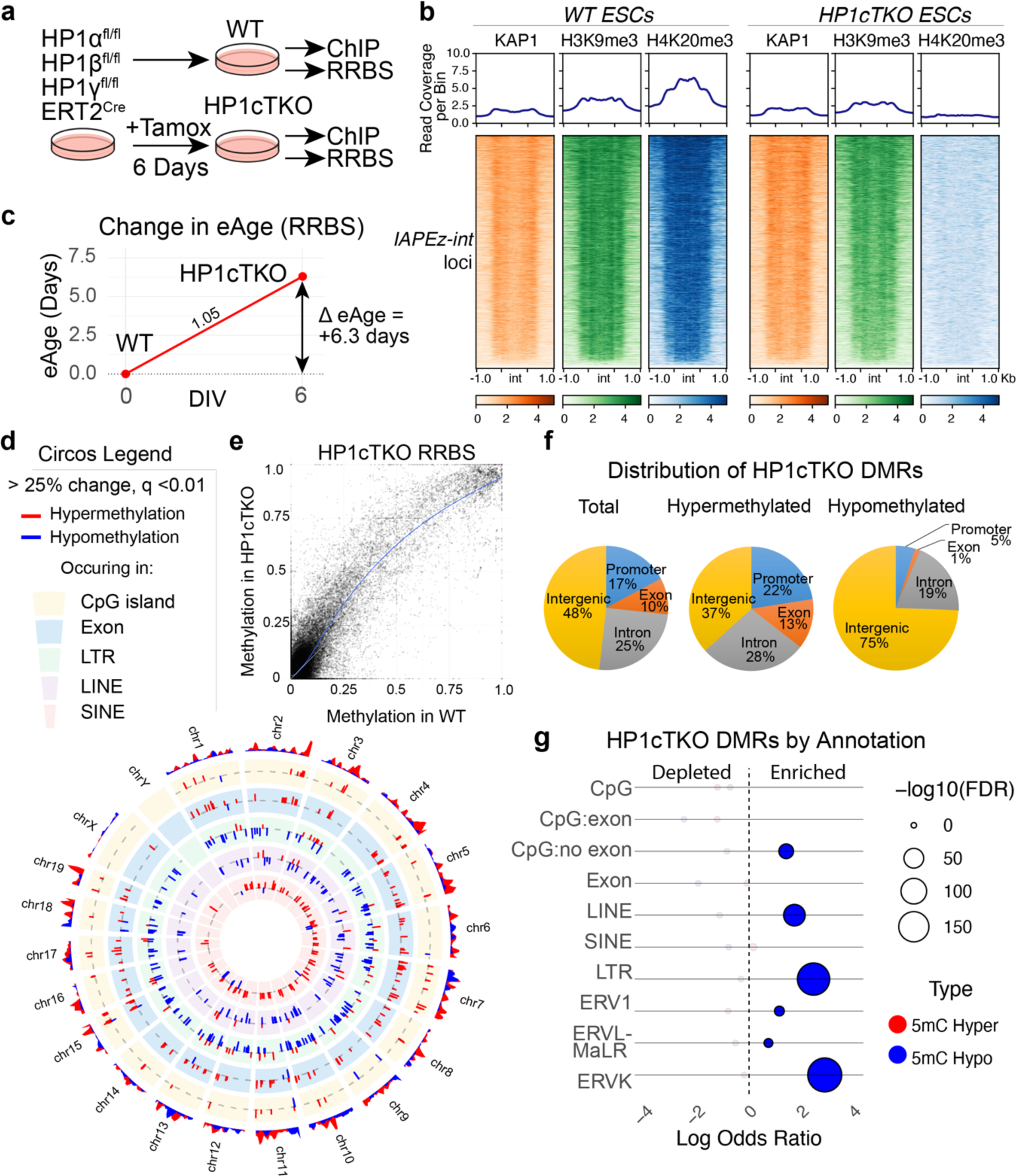
DNA methylation is further perturbed in HP1cTKO ES cells. **a** Schematic of the HP1cTKO RRBS experiment where *HP1α^fl/fl^HP1β^fl/fl^HP1γ^fl/fl^ERT2^Cre^* ES cells are left untreated (WT) or treated with tamoxifen (HP1cTKO) for 6 days in vitro before profiling DNA methylation status by Reduced Representation Bisulphite Sequencing (RRBS) and H3K9me3, H4K20me3 and KAP1 by ChIPseq. **b** Coverage of KAP1, H3K9me3 and H4K20me3 over *IAPEz internal segments (int)* and adjacent LTRs is reduced in HP1cTKO ES cells. Given the deletion of HP1γ in HP1cTKO, H4K20me3 is lost entirely. **c** Profiling of ∼18,000 CpG sites in WT and HP1cTKO ES cells reveals that deletion of HP1 proteins initiates a positive change in eAge. **d** Circos plot of methylation changes that are greater than 25% (q < 0.01) plotted by chromosome by annotation. The magnitude of change is represented on the radial y axis. The outermost ring represents methylation change density. Inner rings annotate methylation changes occurring to CpG islands, exons, LTRs, LINEs and SINEs respectively. **e** Scatterplot of cytosine methylation observed in both HP1cTKO and WT Reduced Representation Bisulfite Sequencing. **f** Distribution of HP1cTKO Differentially Methylated Regions (DMRs) by genic feature. **g** HP1cTKO DMRs by annotation, coloured if statistically significant by q value <0.01 and absolute methylation difference is greater than 25%. (F) Odds ratios of overlap of significant HP1cTKO DMRs plotted against the adjusted p value (FDR) of the respective hypergeometric test.

DNA methylation can be used to accurately estimate the biological or ‘epigenetic’ age (eAge) of tissues, which increases at a steady rate in adult tissues ^7^. While embryonic stem cells normally maintain an assigned eAge of zero ^7^, we were surprised to find that the eAge of HP1cTKO ES cells increased faster than their time in culture (fig. **3c**).

HP1cTKO ESCs displayed hypomethylation at LINEs and LTRs (particularly ERVK) (fig. **3d****,g**) despite a global shift towards hypermethylation (fig.**3e**) which is also represented in promoters and gene bodies (fig **3f**). While a large number of significantly hypermethylated cytosines overlap with CpGs, exons, SINEs (fig **3d**) and ICRs (fig **S5b**), as annotation sets they do not show statistical significance based on the hypergeometric test (Fig. 3g). This lack of significance may be attributed to the global background shift towards hypermethylation.

Given that reduced representation bisulfite sequencing (RRBS) cannot discriminate between 5-methyl cytosine (5mC) and 5-hydroxy-methyl cytosine (5hmC), we surmised that some of the ‘hypermethylation’ observed in the HP1cTKO RRBS samples may be attributable to active demethylation processes, specifically the oxidation of 5mC to 5hmC mediated by TET enzymes.^43^ We tested two specific loci using primers designed to amplify over single HpaII/MspI (CCGG) sites from hippocampal lysates. We found that cytosine methylation at Neuronatin (*Nnat)* is decreased in an age-dependent manner in *HP1β^fl/fl^ Emx1^Cre^* hippocampal lysates. Similarly, *HP1β^fl/fl^HP1γ^fl/fl^Emx1^Cre^* hippocampal lysates showed age-dependent decreases in 5mC methylation at IAP sequences alongside a trend towards hydroxymethylation for both loci (fig. **S6a**).

To elucidate the full effects of HP1 deficiencies on 5mC and 5hmC, we conducted RRBS both with and without prior oxidation on hippocampal lysates. This oxidation step converts 5hmC into 5-formylcytosine (5fC), which does not undergo bisulfite conversion, thereby allowing us to distinguish between 5mC and 5hmC (fig **4a**). After an average 67% unique (+20% ambiguous) read mapping alignment efficiency we found that bisulfite conversion using this method recovered on average ∼35% of Methylated CpGs (fig **4b**). This experiment yielded results largely consistent with what was observed in HP1cTKO ESCs. We found that deficiency of HP1β results in aberrant 5mC (*de novo*) hypermethylation of promoters and CpG islands, a large percentage coming from a gene desert on chromosome 12 (**fig 4c-e, S6c**). Surprisingly, we also found that clustered protocadherin promoters became dramatically 5mC hypomethylated in HP1β/γ DKO hippocampi (fig **4e****, S6d**)., along with imprinting control regions (ZFP57 ICRs) (**fig 4e**). 5mC DNA methylation at LTRs, primarily ERVK elements, was markedly reduced in HP1β/γDKO hippocampi (**Figure 4e, S6e**). This effect is also likely an underestimate given the lower mapability of repeats in RRBS sequencing and a lower bisulfite conversion efficiency in this experiment. We could also observe corresponding statistically significant increases in 5hmC over ERVK and IAPLTR1a annotated regions in HP1β/γDKOs. Notably, single deficiency of either HP1β or HP1γ already initiates drift in DNA methylation, evidenced by significant increases in 5hmC over introns, LINEs, SINEs, and LTRs, an effect that is recapitulated in normal aging in wildtype hippocampi (**fig. 4e**).

**Figure 4.**
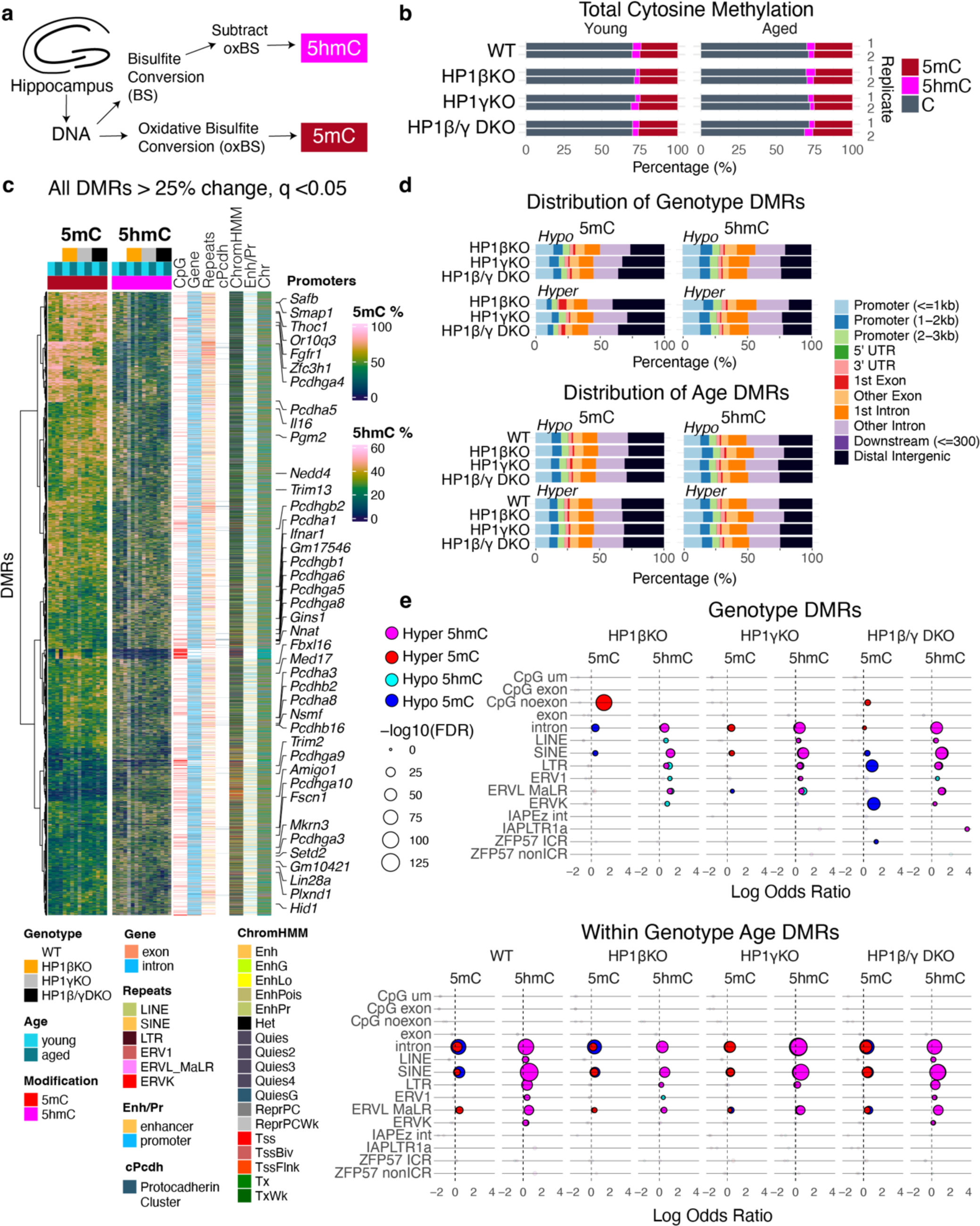
DNA methylation fidelity is progressively compromised in HP1 deficient hippocampi. **a** Schematic of experimental design showing determination of paired 5-methyl Cytosine (5mC) and 5-hydroxy-methyl Cytosine (5hmC) measurements from young and aged *HP1β^fl/fl^;Emx1^Cre^*, *HP1γ^fl/fl^Emx1^Cre^* and *HP1β^fl/fl^HP1γ^fl/fl^Emx1^Cre^* mutant hippocampal lysates by Reduced Representation Bisulfite Sequencing (RRBS). Genotypes that follow are abbreviated to HP1βKO, HP1γKO, and HP1β/γDKO respectively for clarity. **b** Global 5mC and 5hmC methylation across biological replicates sampled. **c** Heatmap of all observed Differentially Methylated Regions (5mC or 5hmC) that change 25% or more and are statistically significant below q = 0.05. DMRs (cytosine positions, rows) observed across genotypes are accompanied by row annotations corresponding to genomic context denoted by being a CpG island, in a gene (intron/exon), repetitive element (LINE, SINE, ERV1, ERVL_MaLR, ERVK or other nonredundant LTRs), overlap with the protocadherin cluster (denoted cPcdh), its chromatin state defined by the 18 state ChromHMM model from P0 mouse cortex (ChromHMM), its overlap with promoters or enhancers defined by the Enhancer-gene map from ENCODE 3, and chromosome (Chr). **d** Total distribution of DMRs by genotype and direction of change (hypomethylation ‘hypo’ or hypermethylation ‘hyper’). **e** Odds Ratios of DMRs significantly changed due to genotype or age are plotted against the q value (FDR) resulting from the hypergeometric test of DMRs overlapping with the annotation. CpG um = Unmasked CpGs, CpG exon = CpG islands overlapping with exons, CpG noexon = CpG islands not overlapping with exons, LTR annotation here refers to all LTRs including ERV1, ERVL MaLR, ERVK etc.

### Activation of astrocytes and microglia

We found that 15% of (88/565 protein coding) genes differentially expressed in *HP1β^fl/fl^HP1γ^fl/fl^Emx1^Cre^* hippocampi overlapped with cellular responses to interferons including *Ifitm2*, *Ifi27* and the regulatory component of the interferon gamma receptor *Ifngr2* (Data S1). HP1DKO hippocampi also showed increased *Oas3*, *Lyk6*, *Wdfy4*, and *Il34*, but most notably displayed elevated transcription of complement *C3, C4b* and *C1qa*.

Given the recently established importance of complement proteins in the developing brain ^44^, their co-occurrence with amyloid plaques ^45^, and their accumulation over normal aging ^30,46^, we performed a multiplex in situ hybridization in order to identify the cell type(s) containing raised levels of Complement 3 (*C3*) RNA (fig. **5a****-i**). Elevated *C3* transcripts could be observed in the soma of wildtype CA1 and CA3 neurons, but in aged HP1β/γ DKO, *C3* could also be detected in small plaque-like foci in the *stratum radiatum* (white arrows, fig. **5d****,g**) that were not observed in any other condition. These foci were surrounded by Iba1+ microglia with a distinct morphology (fig. **5e****,f**). A second experiment revealed these *C3+* foci were also *Slc1a3+* indicating these foci were reactive astrocytes (fig. **5g****-i** & **S7a,b**). In HP1β/γ DKO hippocampi, the number of total GFAP+ astrocytes significantly increases compared to young DKO animals, although this puts it in the same range as the other genotypes. Notably, ∼50% more Iba1+ microglia can be observed in aged HP1β/γ DKO hippocampi (fig. **5j****,k** & **S7c,d**). We could observe the Iba1+ microglia neighboring GFAP+ astrocytic foci in the *stratum radiatum* of HP1DKO hippocampi exhibited large CD68+ protrusions (arrows fig. **5l**), indicating augmented phagocytosis. We quantified the CD68+ area within Iba1+ cells and found that this significantly increased in HP1β/γ DKO hippocampi in an age-dependent manner (fig. **5m**). This suggested that pro-inflammatory signalling, likely as a result of the de-repression of IAPs and other ERVs, chronically results in greater activation of microglia.

**Figure 5.**
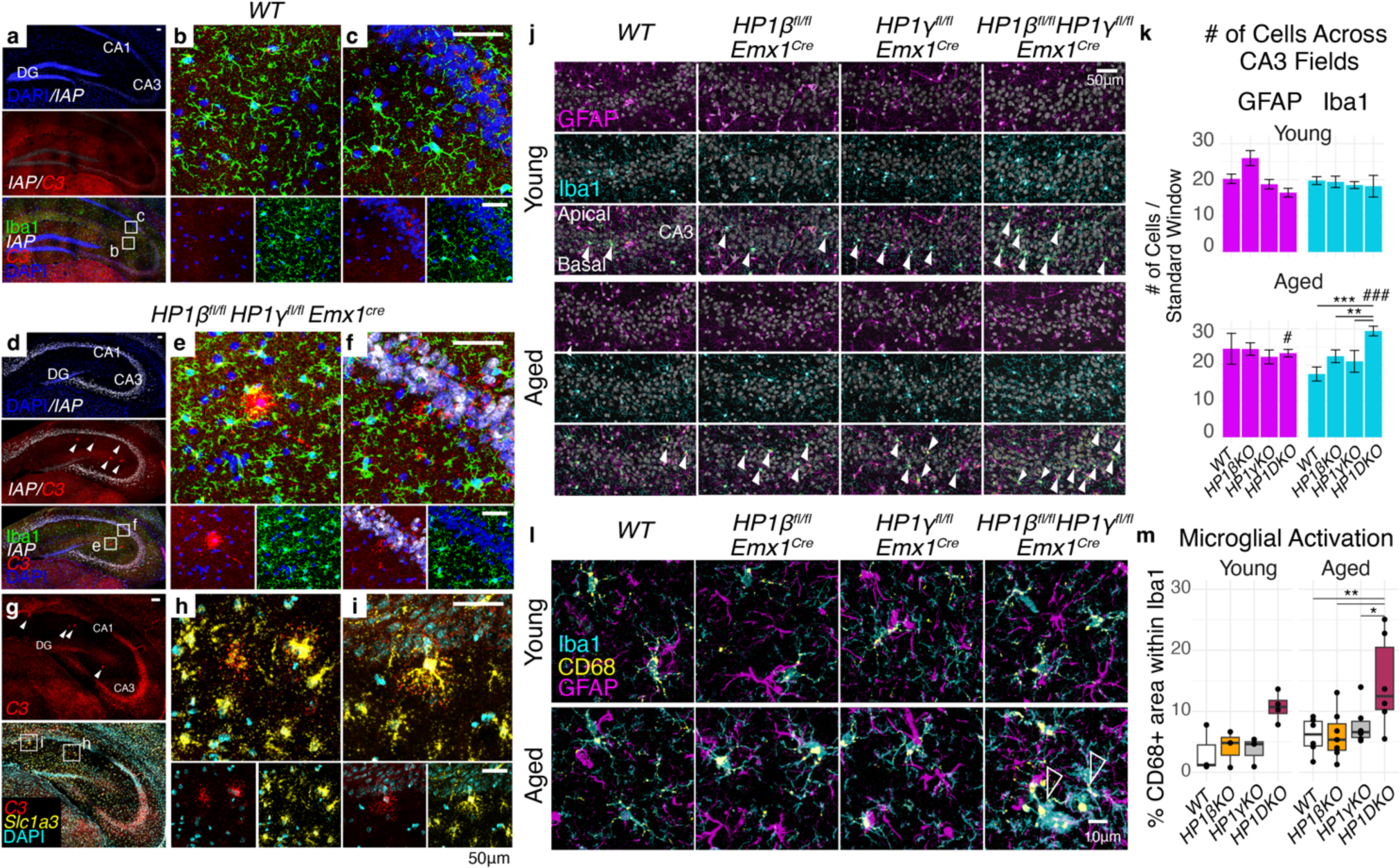
Chronic de-repression of ERVs coincides with the appearance of C3+ reactive Astrocytes and increased CD68+ Microglia. Moderate Complement 3 (C3) RNA can be detected by RNAscope in aged WT **a-c**, where it can be detected in most cells of the hippocampus including CA1 and CA3 pyramids. (Representative images are displayed here after testing across 3 brains per condition). De-repression of IAP transcripts in aged *HP1β^fl/fl^HP1γ^fl/fl^Emx1^Cre^* (white, **d**) corresponds with large C3+ islands (arrows in **d**) that can be found in the *Stratum Radiatum* (magnified in **e**) and emanating from the *Stratum Pyramidale* (magnified in F). Such C3 foci are surrounded by microglia with distinct morphology (compare **b** & **c** to **e** & **f**). C3+ foci found in the *Stratum Radiatum* of *HP1β^fl/fl^HP1γ^fl/fl^Emx1^Cre^* hippocampi are Slc1a3+ astrocytes (**g**, magnified in **h** & **i**). **j** Representative images of CA3 pyramidal layers stained for GFAP, Iba1 and Dapi. Iba1+ cells entering the *stratum pyramidale* are indicated with solid arrows. **k** Quantification of GFAP+ and Iba1+ cells across apical, somal, and basal regions in CA1 and CA3 fields (see also fig S7). Statistics two-way ANOVA with estimated marginal means post-hoc test with Tukey’s familywise correction. Young: WT n= 8, HP1βKO n = 10, HP1γKO n=8, HP1DKO n=10, Aged WT n=8, HP1βKO n= 14, HP1γKO n=8, HP1DKO n= 14. Adjusted p values for genotype test within age (Iba1): aged HP1DKO vs WT p=0.003, aged HP1DKO vs aged HP1γKO p=0.0152, aged HP1DKO vs aged HP1βKO p = 0.0152. Adjusted p values for age test within genotype: Iba1 HP1DKO age p<0.0001, GFAP HP1DKO age p = 0.0087. **l** Iba1+ microglia that can be found in the *stratum radiatum* surrounding GFAP+ astrocytes contain large CD68+ compartments suggesting endosomal activity. **m** Quantification of Microglial activation (from **l**) measured by the proportion of CD68+ area within Iba1+ microglia. Statistics two-way ANOVA with estimated marginal means post-hoc test with Tukey’s familywise correction. Young: WT n= 3, HP1βKO n = 3, HP1γKO n=3, HP1DKO n=4, Aged WT n=6, HP1βKO n= 7, HP1γKO n=6, HP1DKO n= 6. Adjusted p values for genotype test within age: aged HP1DKO vs WT p=0.008, aged HP1DKO vs aged HP1γKO p= 0.0444, aged HP1DKO vs aged HP1βKO p = 0.0064.

While it is known that proteins derived from ERVs can be pro inflammatory in the brain^47^, the majority of ERV transcripts do not have complete coding sequences. Given that IAP transcripts de-repressed in *HP1β^fl/fl^/HP1γ^fl/fl^Emx1^Cre^* brains are neuronally derived and glial activation follows, we designed an experiment to profile the stimulatory effect of IAP ssRNA on mixed glial cultures *in vitro* (fig **6a**). As a control, we used the same sequence derived from IAP but substituted pseudo uracil^48^, thus termed ψ-IAP (fig **6b**). Twenty-four hours following introduction of IAP ssRNA we profiled the media and cytosolic lysate for activation of chemokines and cytokines using a membrane-based sandwich immunoassay (fig **6c**). We found robust activation of several classical cytokines in response to IAP ssRNA including TNF, IL-6, IL-12, IL-16, IL-23 (fig **6d**). The most prominent activation was in CCL3, CCL4 and CCL5 (RANTES). We confirmed CCL5 activation predominantly came from GFAP+ astrocytes (fig **6e**) and this was specific to IAP ssRNA and not ψ-IAP ssRNA (fig **6f**). Interestingly, while IFN-γ increases were not statistically significant in our assay, the cytokine response to IAP ssRNA observed here are consistent with previous observations of IFN-γ stimulated increases of ERV dsRNA, which includes elevated IL-6 and CCL5^49^.

**Figure 6.**
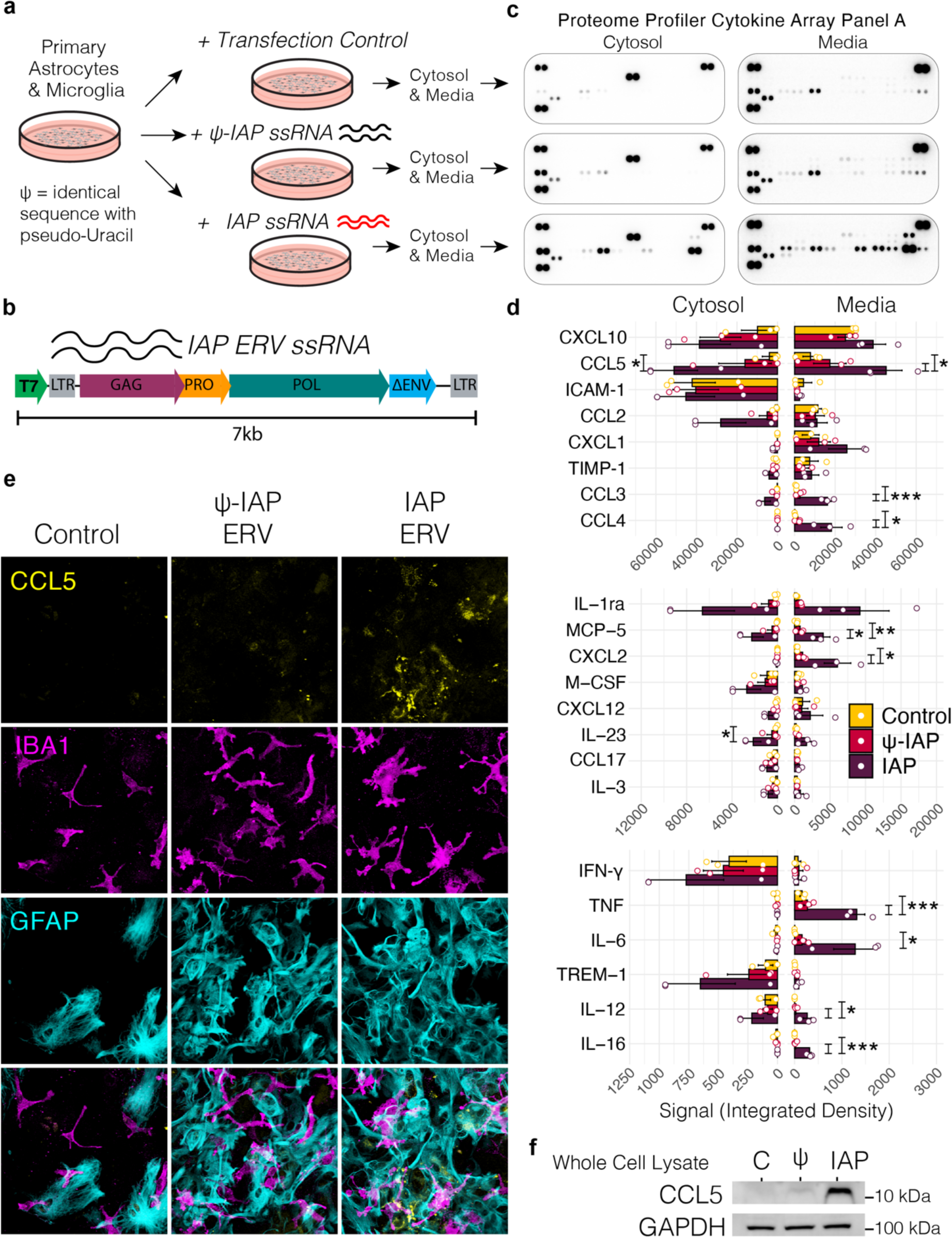
Acute introduction of IAP ssRNA induces an inflammatory response. **a** Schematic showing experimental design where primary astrocytes and microglia are exposed to a pulse of regular- or pseudo (ψ)-IAP ssRNA, where ψ-IAP is generated from the same template but with ψ-uracil. **b** Full length IAP template used for generation of ssRNA. **c** Representative dot plots from Proteome Cytokine Array Panel A following incubation with media or cytosolic lysates from control, ψ-IAP ssRNA or IAP ssRNA treated glial cultures. **d** Quantification of Cytokine Panel A immunoassays, performed in 3 biological replicates separated by cytosol and media, split by high signal (top), medium signal (middle), and low signal (bottom). **e** Immunofluorescence stain of glial culture comprised of Iba1+ microglia and GFAP+ astrocytes stained for CCL5 (RANTES) and GAPDH loading control following incubation with ψ-IAP and IAP ssRNA.

### Increased Dendritic loss and Cognitive Decline

Young HP1β KO CA3 hippocampal neurons showed a modest reduction in CA3 basal dendrite complexity that was carried through to aged animals. By contrast, HP1β/γ DKO animals displayed a pronounced age-dependent decrease in CA3 basal dendrite complexity (fig. **7a**). Structural MRI revealed that while young HP1β/γ DKO animals have a modestly reduced volume of both the cortex and total brain, this difference is less obvious in aged HP1β/γ DKO animals (fig. **S8**).

**Figure 7.**
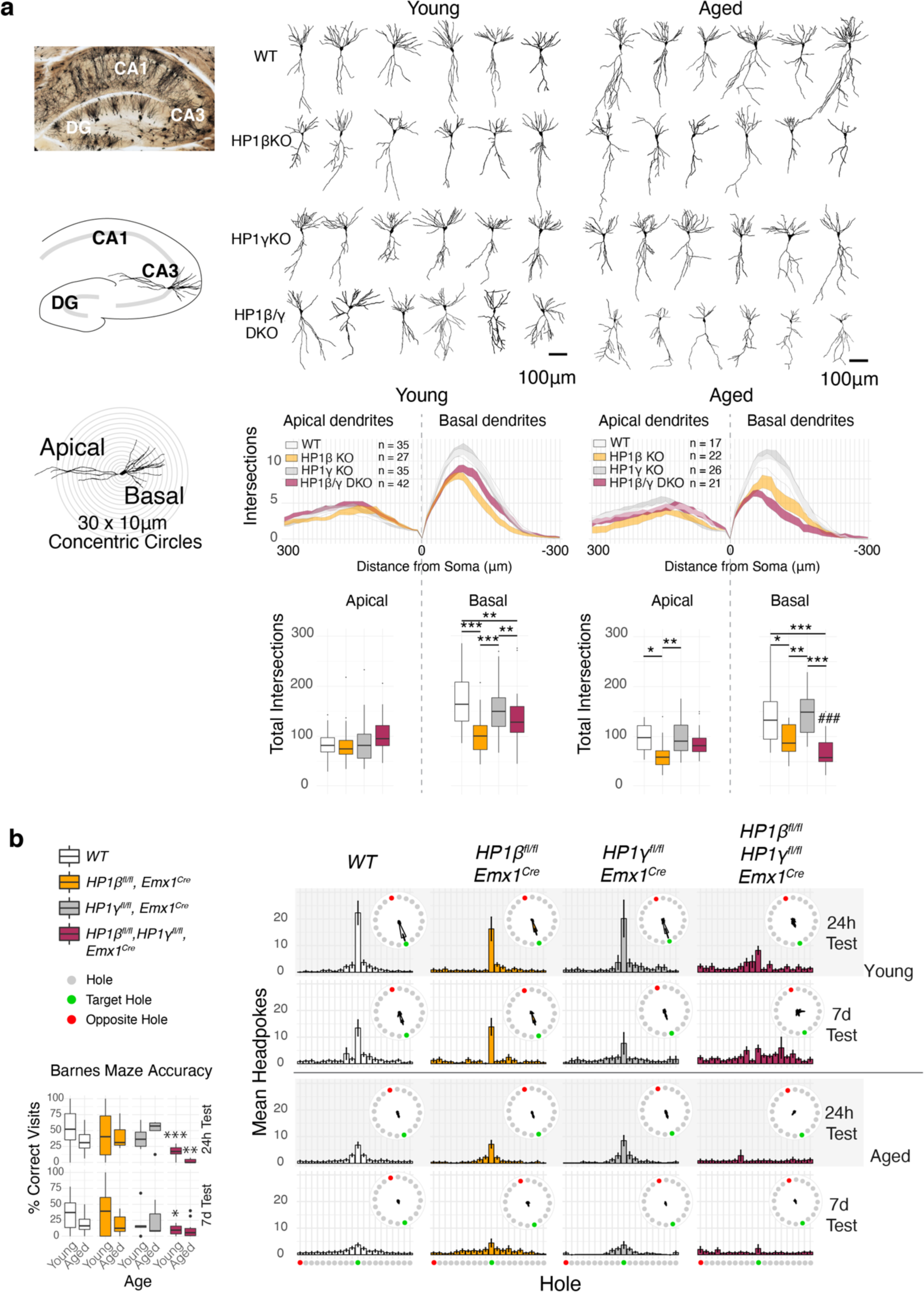
HP1 deficiency causes age related behavioral abnormalities and CA3 dendritic tree degeneration. **a** CA3 Dendritic Complexity in young and aged HP1 mutants by Golgi impregnation and Scholl analysis. Deficits in basal dendrite complexity can be observed in young HP1βKO and HP1β/γ DKO animals. While basal dendrite complexity is nearly identical in aged HP1βKO, HP1β/γ DKO basal dendrites show an age dependent degeneration, losing almost 50% of their complexity. **b** Histograms of performance in the circular Barnes Maze (green = target hole, red = opposite). Histograms plot mean headpokes ±SEM. Asterisks mark ANOVA across genotypes within age, *** p < 0.001, ** p < 0.01, * p < 0.05. Hashtags mark ANOVA across age within genotype, ### p < 0.001.

Cognition and behavior of HP1 mutant mice was profiled as young adults (3-4 months) and again in middle age (13-14 months), where we observed several age-dependent deficits. HP1β/γ DKO animals exhibited deficits in spatial learning and memory as tested in the Barnes maze (fig. **7b**). Young HP1β/γ DKO animals showed impairment in both learning (24h test) and recall (7 day test) of the target nest, while aged HP1β/γ DKO animals seemed unable to learn the location of the target nest and typically walked around the periphery continuously. Aged HP1β/γ DKO animals also showed an age-dependent abolition of paired-pulse inhibition (fig. **S9a**), extended bouts of eating and grooming (fig **S9b**) and an altered circadian rhythm (fig. **S9f**). During handling and testing, a subset of HP1 mutant animals displayed stimulus-dependent seizures, which were observed with HP1β KO animals predominantly (fig. **S9e**). Young HP1β/γ DKO animals displayed hyperactivity in an open field, and as aged animals they showed both an absence of center zone anxiety (fig. **S9c**) and a marked inability for nest construction (fig. **S9d**).

## Discussion

Our study has uncovered nonredundant functions of HP1β and HP1γ in neurons. The age-dependent reduction in DNA methylation at imprinted loci as observed for *Nnat* in *HP1β^fl/fl^Emx1^Cre^* indicates that HP1β stimulates Dnmt1 activity, as has been described previously^50^. That the two proteins can operate along the same pathway would explain the similarity of the reduction in the formation of the dentate gyrus in tamoxifen induced *Nestin-Cre* deletion of *Dnmt1*^51^ compared to that observed here with *HP1β^fl/fl^Emx1^Cre^*. Moreover, the robust activation of IAP elements in *Dnmt1^fl/fl^Emx1^Cre^* mutant phenocopies the *HP1β^fl/fl^HP1γ^fl/fl^Emx1^Cre^* mutant, albeit the *Dnmt1^fl/fl^Emx1^Cre^* mutant also displays a dramatic malformation of the cortex ^52^, which we did not observe. Notably, many of the genes upregulated in *HP1β^fl/fl^HP1γ^fl/fl^Emx1^Cre^* hippocampi overlap with regions where Dnmt1-dependent methylation is not recovered once abolished ^53^. Since 5mC methylation is dramatically lost at ERVK elements in *HP1β^fl/fl^HP1γ^fl/fl^Emx1^Cre^* but not in single mutants, this suggests that *HP1β or HP1γ* are capable of stimulating Dnmt1 activity and the loss of both is detrimental. Age-related 5mC hypomethylation appears to be restricted to loci already under threat of activation such as the tissue-specific imprinted genes (*i.e., Nnat*) or partially silenced ERVK repeats such as IAPs. This phenomena has been observed before with IAP elements in aging mice—where their periodic activation results in progressive demethylation and complete de-silencing ^54^, and may explain why IAP transcription further increases in aged *HP1β^fl/fl^HP1γ^fl/fl^Emx1^Cre^* (fig. **1c**). The shift towards hydroxymethylation in *HP1β^fl/fl^HP1γ^fl/fl^Emx1^Cre^* hippocampi along with the measured increases in age-related hydroxymethylation in non-coding regions (fig **4e**) indicate that HP1 proteins also serve to protect against TET-mediated hydroxymethylation.

*HP1β^fl/fl^HP1γ^fl/fl^Emx1^Cre^* hippocampi show several functional similarities with the transcriptional signatures of very aged (24-29 month) mouse hippocampi. Very aged hippocampi display downregulation of ZFP57, Suv420h2, and upregulation of much of the protocadherin cluster ^55^. Some of the strongest transcriptional changes in *HP1β^fl/fl^HP1γ^fl/fl^Emx1^Cre^* hippocampi are also the strongest observed changes in very aged hippocampi, including age-associated ncRNAs *Pisd-ps1* and *Pisd-ps2* and complement components *C1qa, C3*, and *C4b* ^46,55–57^.

*HP1β^fl/fl^HP1γ^fl/fl^Emx1^Cre^* hippocampi also show similarities to the ‘pre-plaque’ state^58^. According to current concepts, the mechanism by which astrocytes and microglia co-ordinate the complement cascade involves the initial_activation of microglia. Activated microglia then secrete Il-1a, TNF and C1q that induce A1-astrocyte reactivity ^59^. In addition to being directly neurotoxic ^59^, it is thought reactive astrocytes also greatly enhance the susceptibility of aging brains to neurodegeneration because they are major source of classical complement cascade components C3 and C4b that then drive microglia-mediated synapse loss ^58^. In Alzheimer’s disease, microglial activation and synapse loss precedes plaque pathology in the hippocampus ^58^, and reactive astrocytes have long been associated with senile amyloid plaques (for review see ^60^).

Although it is unclear how microglia are activated in *HP1β^fl/fl^HP1γ^fl/fl^Emx1^Cre^* hippocampi, a likely sequence of events is that microglia activation is a response to the export of ERV or other inflammatory RNA from *HP1β^fl/fl^HP1γ^fl/fl^Emx1^Cre^* neurons that is detected by astrocytes and microglia. Similar export of RNA or protein-aggregate filled extracellular vesicles (EVs) have been observed in neuron-neuron or neuron-glia communication ^61,62^, and ERVs have been shown to be included in EVs ^63^. In this context, human ERVs have been found to be elevated in Alzheimer’s ^17^ and ALS ^20^. Extracellular vesicles that export neuronal unfolded protein aggregates have also been shown to serve as an activating signal to microglia or astrocytes ^62^ and this is sufficient to drive non-cell-autonomous neuronal degeneration ^64^.

When we tested the effect of extracellular ERV RNA on mixed glial cultures, we could observe an inflammatory response akin to direct stimulation of IFN-γ ^49^, which is also consistent with the upregulation of a large number of interferon related genes we observed in RNaseq (fig. **1**). We also note the upregulation of IFNGR2, the regulatory subunit^65^ of the interferon gamma receptor in *HP1β^fl/fl^HP1γ^fl/fl^Emx1^Cre^* hippocampi. Activated astrocytes in many neurodegenerative diseases show upregulation of IFNGRs^66^, suggesting an increase in sensitivity to IFN-γ signaling. Once effected, IFN-γ signaling has been observed to up-regulate expression of the complement components C3 and C4 by stabilization of their mRNA ^67,68^. Notably, in addition to other non-inflammatory roles, IFN-γ signaling has been shown to promote tau hyperphosphorylation ^69^.

The extensive cross-talk that occurs between innate immune pathways and the unfolded protein response (UPR) is known collectively as the integrated stress response (ISR) ^70^. UPR engagement in *HP1β^fl/fl^HP1γ^fl/fl^Emx1^Cre^* hippocampi is likely a direct result of sustained ERV transcription. It is known that extended UPR stress in astrocytes results in Complement Activation^64^. Our results are also consistent with observations that TDP-43 proteinopathies, which can be potentiated by elevated ERV expression^22^, also result in activation of complement^71^. Given the majority of neurodegenerative diseases are characterized by misfolded proteins or altered proteostasis ^70,72^, this provides an invaluable insight into how heterochromatin loss alone can drive core components of neurodegeneration^73^ and accelerated cognitive decline.

Complement activation in the aging brain may have as many exogenous inducers as there are pathways to inflammation. However, the present study indicates that at least one primary trigger of the innate immune response is of endogenous origin: namely, the activation of endogenous retroviruses following heterochromatin loss.

## Acknowledgments

Computation was performed on the HPC for Research/Clinic cluster of the Berlin Institute of Health. We would like to acknowledge Julie Brind’Amour, Carol Chen and Matthew Lorincz for early discussions. We thank Marion Rivalan and Melissa Long from the NeuroCure Animal Outcome Core Facility (AOCF) for facilitating behavioral experiments. We thank Robert Schwarz for statistical advice on calculating Odds Ratios. We also thank Seija Lehnardt and Christina Krüger for advice with astrocyte culture. We would also like to thank Ingo Bormuth, Roman Wunderlich, Ulrike Günther, Denis Lajkó and the animal facility of the Charité.

## Author contributions

Conceptualization, A.G.N, P.B.S and V.T.; Investigation, A.G.N., J.S., S.Z., P.B., S.M., P.B-S.; Resources, A.G.N., J.B., M.N., O.O., H.K.; Bioinformatics, A.G.N.; Formal analysis, A.G.N.; Visualization, A.G.N.; Funding acquisition, A.G.N., P.B.S., V.T.; Writing-original draft, A.G.N., Writing-review and editing, A.G.N., P.B., P.B.S., V.T., Writing-revision A.G.N.; All authors reviewed the final manuscript.

## Data availability

Data from this study has been deposited at GEO under accession # GSE153331

## Code availability

Code used in analysis can be accessed at https://github.com/qoldt/HP1-Deficiency-Neurodegeneration

## Methods

### Mouse Lines

All animal experiments were conducted in compliance with animal welfare guidelines put in place by the für Gedundheit und Soziales (LaGeSo) Berlin.

The **H**eterochromatin **P**rotein **1 F**loxed **E**mx1**C**re (HP1FEC) mouse line was generated by generating floxed alleles of HP1β (*Cbx1*) and HP1γ (*Cbx3*) which were combined with the Emx1-IRES-Cre mouse (Jackson Laboratory). The HP1β (*Cbx1*) targeted allele was generated by introduction of a targeting cassette into ES cells by homologous recombination that inserted loxP sites surrounding exons II and III. Successful integrations were detected by neomycin selection via an FRT flanked neomycin cassette inserted between exons III and IV. Following a cross to a flip-deleter mouse, the NeO cassette is removed by flippase-mediated recombination, giving rise to the *HP1β^fl/fl^* allele (Figure S10). HP1γ (*Cbx3*) was targeted using a similar strategy giving rise to the HP1γ^fl/fl^ allele (Figure S11).

### HEK Cell Culture & Transfection

HEK293T cells were cultured in DMEM+Glutamax (Gibco) supplemented with 10% FBS and 1% penicillin/streptomycin (Gibco). For Co-IP experiments, HEK293T cells were seeded in 6 well plates at a density of 0.3 x 10^6^ cells per well. For IHC, cells were plated onto glass coverslips in 24 well plates at 100,000 cells per well. Plasmid DNA was introduced into cells by chemical transfection using Lipofectamine 2000 according to the manufacturer’s instructions.

### Co-Immunoprecipitation (Co-IP) and Western Blot (WB)

Six well plates were rinsed briefly in cold PBS, then lysed on ice in 300μl of Radioimmunoprecipitation assay buffer (RIPA, 50 mM Tris, pH 8.0, 150 mM NaCl, 1% Triton X-100, 0.5% Sodium Deoxycholate, 0.1% SDS) supplemented with 1X Protease Inhibitor Cocktail (PIC, Roche). Lysates were then sonicated for 15 pulses on ice using a probe sonicator. Insoluble debris was then precipitated by centrifugation at 13,000 rpm at 4°C, and then decanted into a new tube.

Protein concentration was measured using a standard Bicinchroninic acid (BCA) assay, by using 10μl of protein sample diluted 1:10, plating 25μl into a 96 well plate in triplicate along with a Bovine Serum Albumin (BSA) standard (0, 0.1, 0.2, 0.5, 0.75, 1.0, 1.5, 2.0 μg/μl).

Prior to immunoprecipitation, each sample was adjusted to a 300μl volume at a protein concentration of 1.5 μg/μl using lysis buffer, and 20μl of lysate was set aside for input. To perform the immunoprecipitation, 1.5μl of antibody (mouse anti-myc 9B11, Cell Signaling or goat anti-GFP, Rockland) was incubated per sample on a rocker for 2 hours at 4°C. During this time, protein G sepharose beads (GE Healthcare, 25 μμl per sample) were rinsed 3 x 15 minutes in 1ml of cold TBS (50 mM Tris pH 7.5, 150 mM NaCl), rocking at 4°C. Between washes beads were spun down using a short 5 second spin on a tabletop centrifuge (no greater than 9,000 rpm). Following the 2 hour antibody/lysate incubation, washed beads are added to each sample and incubated on a rocker for an additional hour at 4°C. Following IP, beads are washed twice using lysis buffer and twice with TBS. On the final wash, as much buffer was removed as possible before addition of 25 μμl 2.5X lammeli buffer. IP was then boiled at 95°C for 5 minutes. For input samples, 5 μμg of total lysate was used in a volume of 25 μμl, adjusted with 5 μμl 5X lammeli buffer and the appropriate amount of lysis buffer, and boiled at 95°C for 5 minutes.

### RNA Isolation

For cDNA library construction, RNA was purified from P0 cerebral cortices using TRIzol (Invitrogen) according to the manufacturer’s instructions followed by reverse transcription by SuperScript II (Thermo Fischer) using random primers. For RNAseq, RNA was isolated using the Relia-prep RNA mini kit (Promega) according to the manufacturer’s instructions.

### RNAseq Library Preparation

RNA-seq libraries were prepared with the NEBNext Ultra RNA Library Prep Kit for Illumina (New England Biolabs), using 1 μg total RNA per experiment.

### Molecular Cloning

For in situ probes, primers were designed around consensus sequences obtained from repbase (https://www.girinst.org/repbase/) and amplicons were queried using UCSC’s BLAT and in silico PCR tools to determine the estimated diversity of transcripts corresponding to the probe. Probe primers for a unique IAP element on chr 2 were inferred based on unpublished qPCR primers (Julie Brind’Amour, UBC) (Data S1). Probe sequences were amplified from cDNA using GoTaq Polymerase (Promega) and ligated into the pGEM®-T vector (Promega). Linearized plasmids were then used as templates for in vitro transcription using either SP6 or T7 (Roche) using DIG labelled nucleotides (Roche). Following DNA digestion and RNA purification, the RNA probe was resuspended in 20μl water and 180μl hybmix (50% formamide, 5X SSC pH 7.0, 1% Boehringer block, 5mM EDTA, 0.1% Tween-20, 0.1% CHAPS, 0.1 mg/ml Heparin, 100 μμg/ml Yeast tRNA) and stored at -20°C until needed.

To clone HP1 and SUV420H2 expression constructs, two rounds of PCR were conducted using the high fidelity Q5 Polymerase (New England Biolabs). The first round of PCR amplified the ‘naked’ coding region of the gene (*_110 and *_111 primers), and using this product as a template, a second PCR was performed with primers removing the stop codon and containing restriction sites. These amplicons were then A-tailed using GoTaq polymerase (Promega) and ligated into the pGEM®-T vector (Promega) yielding pGEMT-KnpI-HP1x (where x is α, β, or γ). To generate eGFP fusions, inserts were digested with KpnI and AgeI for insertion into pCAG-eGFP (Clontech), yielding pCAG-HP1x-GFP fusion constructs. Prior to cloning SUV420H2, pCAG-mycDKK was created by substituting the CAG promoter from pCAG-eGFP with the CMV promoter of pCMV6-Entry though digestion with SpeI and EcoRI. First, ‘naked’ SUV420H2 was amplified using 110 and 111 primers, then 1μl of this template was used for a PCR using kpnI and XhoI primers. The KpnI-SUV420H2-XhoI PCR product was then gel-extracted and digested for 20 minutes using Fast Digest KpnI and XhoI (Fermentas) and subsequently ligated into KnpI/XhoI digested pCAG-mycDKK, resulting in an in-frame C-terminal myc-DKK (Myc tag and Flag tag).

For SUV420H2/HP1 IP experiments, mutations were created in the chromoshadow domain of HP1γ in pGEMT-KpnI-HP1γ using the Q5 site-directed mutagenesis kit (New England Biolabs) according to manufacturer’s instructions (Data S1). Following sequence verification, mutant HP1γ inserts were ligated into pCAG-eGFP as above. Expression plasmids from this study have been deposited at https://www.addgene.org/browse/article/28223393/.

For cloning of the full length IAP construct for in vitro transcription, pFL, a plasmid encoding full length IAP ^74^ and kind gift of Prof. Horie, was first subcloned to delete the antisense intronic GFP using a NEB KLD reaction, and then a T7 promoter sequence was added 5’ to the LTR by NEBuilder gateway cloning, resulting in the plasmid pT7-IAP. Primers can be seen in Data S1.

### In vitro transcription of IAP ssRNA

The pT7-IAP plasmid was linearized using NaeI and NotI for 4 hours at 37°C using 15ug of plasmid DNA. The linearized 6.8kb fragment was then gel purified & used as a template for in vitro transcription (IVT) using the RiboMax large scale RNA production system (Promega, cat # P1300) with minor modifications. First, 3’ overhangs were filled in by incubating 10ug of DNA template with DNA polymerase I large (Klenow) Fragment in T7 transcription buffer for 15 minutes at 22°C. Then T7 in vitro transcriptions were set up with rNTP mixes. For IAP RNA a normal mix of rNTPs (100mM ATP, 100mM UTP, 100mM CTP, 100mM GTP), and for pseudo-IAP, pseudo-uracil was substituted (100mM ATP, 100mM Ψ-UTP, 100mM CTP, 100mM GTP). IVT reactions were then incubated with T7 & respective rNTP mixes for 4hrs at 37°C. Following in vitro transcription, RQ1 RNase-free DNase was added at a concentration of 1u per μg of template DNA and incubated for 15 minutes at 37°C. Then, RNA was extracted in the aqueous phase after addition of 1 volume of citrate-saturated phenol (pH 4.7):chloroform:isoamyl alchohol (125:24:1) (Sigma, cat # 77619), and a second addition of 1 volume of chloroform:isoamyl alcohol (24:1). The aqueous phase was then transferred NAP-5 (GE Healthcare Cat# 17-0853-01) chromatography columns for the removal of unincorporated ribonucleotides and eluted using ultrapure RNase free water. RNA was then precipitated by addition of 0.1 volume of 3M Sodium Acetate (pH 5.2) and 1 volume of isopropyl alcohol, mixing well and incubating 5 minutes on ice. Precipiated RNA was then pelleted by centrifugation at 13,000 in a tabletop centrifuge at 4°C. Supernatant was then discarded and the RNA pellet washed in 70% ultrapure EtOH, dried & resuspended in a volume of Ultrapure ddH2O equal to the transcription reaction. RNA concentration was then measured by first diluting 2μl of RNA into 298μl of water and measuring the absorbance at 260nm. The concentration of RNA was then calculated using the expression

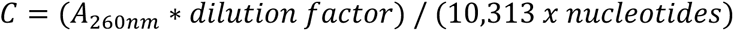

where *C* is in moles and the dilution factor is 100. In vitro transcribed RNA was then confirmed by denaturing 1μg of RNA at 65°C for 10 minutes in 1.5X probe buffer (60% formamide, 40% glycerine 6% formaldehyde, 5% ethidium bromide, 5% bromophenol blue, 20mM MOPS, 5mM EDTA, 2.1mM Calcium Acetate) and running on a 1% agarose gel (20mM MOPS, 5mM EDTA, 2.1mM Calcium Acetate, 6% formaldehyde, 1 % agarose) using MOPS (20mM MOPS, 5mM EDTA, 2.1mM Calcium Acetate) as the running buffer.

### Transfection of Primary Astrocytes & Microglia for Cytokine Profiling

Astrocytes and microglia were obtained by trypsin-assisted dissociation of P0 cortices and cultured for two weeks in DMEM supplemented with 10% FBS and 1% penicillin/streptomycin. Mixed glia were then seeded on poly-D-lysine coated plates at 500,000 cells per cell of a six well plate and 100,000 cells per well of a 24 well plate. For one biological replicate 4 wells of a six well plate per pooled per experiment. RNA was put into complex with lyovec (Invivogen, cat # lyec-1) to a final concentration of 10ng/μl, and 100μl (1μg RNA) applied per six well and 25μl (250ng RNA) applied per 24 well. After a twenty-four hour incubation, conditioned media was removed and cells were rinsed in cold PBS before being lysed in 300μl (across 4 wells of a six well plate) using cytokine lysis buffer (1% Igepal CA-630, 20 mM Tris-HCl (pH 8.0), 137 mM NaCl, 10% glycerol, 2 mM EDTA, 10 μg/mL Aprotinin, 10 μg/mL Leupeptin, and 10 μg/mL Pepstatin). Cells were lysed for 30 mins on ice with gentle pipetting. Proteome Profiler Cytokine Array Panel A (R&D Systems, cat # ARY006) was then used for parallel detection of activated cytokines and chemokines in cytosolic lysates or conditioned media according to manufacturer’s instructions. Quantification of dot plots was performed using the Quick Spots software (Ideal Eyes Systems, Inc.).

### In situ Hybridization

Standard DIG-labelled in situ hybridization was performed using the method described in^75^. Multiplex in situ hybridization was performed using RNAscope (Advanced Cell Diagnostics) (*3*, *4*) (*3*) according to the manufacturer’s instructions using a custom designed probe for IAP and standard C3 and Slc1a3 probes. Iba1+ cells were identified after RNAscope using an Iba1 antibody (Wako).

### In utero electroporation

In utero electroporation was performed according to the initial protocol ^76^ with minor modifications. All surgical procedures were performed in accordance with LaGeSo experimental licenses G0079/11 and G0206/16.

### Tissue Processing & Histology

For embryonic tissue, the date of the vaginal plug was counted as embryonic day 0.5 (E0.5). Pregnant females at the desired stage were killed by an i.p. injection of 600 mg pentobarbital per kg body weight and death was confirmed by cervical dislocation. Following a midline incision, uterine horns were excised and placed in a dish containing ice cold PBS, where embryonic brains were isolated using microforceps. Brains were immediately fixed in paraformaldehyde (PFA, 4% in PBS) overnight at 4°C. For animals older than P6, animals were given a lethal i.p. injection of 600 mg pentobarbital per kg body weight and transcardially perfused with PBS (10-20ml, depending on age) until the liver was clear, followed by perfusion with 4% PFA (5-20ml, depending on age). Brains were then isolated and fixed overnight in 4% PFA.

For in situ hybridization, all solutions were composed using ddH_2_O treated with Diethyl pyrocarbonate (DEPC, Sigma, prepared by shaking 1ml DEPC in 1L ddH_2_O at 37°C O/N) and per processed by cryosectioning. For cryosectioning, after fixation brains were dehydrated in sequential concentrations of sucrose (15%, 30% in PBS) before being embedded in Optimal cutting temperature compound (OCT, Tissue-Tek) and freezing on dry ice. Tissue blocks were stored at -20°C until 16μm sectioning on a cryostat at -20°C, when they were collected on positively charged slides (Superfrost, ThermoFischer) and allowed to dry for 1 hr before being re-frozen for storage at -20°C.

For paraffin sectioning, fixed brains were dehydrated by ethanol row (30% EtOH, 50% EtOH, 70% EtOH, 80% EtOH, 90% EtOH, 100% EtOH) followed two changes of Xylol and two changes of paraplast before casting in paraffin in metal embedding molds. For sectioning, a Microtome (Leica) was used and 14μm thick sections were collected in 37°C ddH_2_O on positively charged slides (Superfrost, ThermoFischer).

Nissl Stains were performed by incubation with cresyl violet. Cresyl violet staining solution was prepared by dissolving 0.1g cresyl violet acetate in 100ml ddH_2_O O/N. Following addition of 10 drops (∼0.3ml) glacial acetic acid, this cresyl acetate solution was filtered. Nissl stain was performed on rehydrated paraffin sections (Xylol II: 5 mins, Xylol I: 5 mins, 3×5 min 100% Ethanol, 3 min 95% Ethanol). Sections were immediately stained in 0.1% Cresyl violet for 3-10 minutes, followed by rinsing in dH_2_O to remove excess stain. Sections were then differentiated in 95% ethanol for 2-30 minutes, checking microscopically for optimal staining. Sections were then dehydrated by 2x 5 min 100% ethanol washes and cleared with xylol (2 x 5 mins) before mounting with Entellan (Sigma).

Immuno Histochemistry (IHC) was performed primarily on cryosections. For paraffin sections, prior to IHC sections were rehydrated and an antigen retrieval step (Boiling 3 x 5 mins in Antigen Unmasking Solution, Vector Labs) was performed prior to blocking. To perform IHC, slides were washed 2 x 5 minutes in PBS, then blocked and permeabilized in blocking solution (2% BSA, 1% Triton X100 in PBS). All further antibody steps use this same blocking solution as diluent. Primary antibodies were diluted 1:200-500 in blocking solution and incubated on sections at 4°C O/N. The following day, slides were washed 3×10 minutes in PBS and then incubated for two hours at room temperature with the appropriate secondary antibody (Dianova). Sections were then washed 2 x 5 mins and stained with Hoechst/DAPI (1:5000 in PBS) for 5 minutes at room temperature. When using adult sections, lipofuscin autofluorescence was quenched by a 10 min incubation with a solution containing 10mM CuSO_4_ & 50mM NH_4_Cl. Sections were then mounted aqueously using Immu-Mount (Shandon). A full list of antibodies used can be seen in Data S6.

For Golgi impregnation, fresh brain samples from WT, HP1β^fl/fl^Emx1^Cre^, HP1γ^fl/fl^Emx1^Cre^ and HP1β/γ^fl/fl^Emx1^Cre^ of young (3 months) and aged (13-14 months) mice were cut into two hemispheres and impregnated in Golgi-Cox solution for 2 weeks as described in ^77^. Sholl analysis ^78^ was performed blind on CA3 hippocampal neurons by using the concentric circles and cell counter plug-ins available for ImageJ. Intersections were quantified across thirty 10μm spaced concentric circles. The Simple neurite tracer plugin (ImageJ) was used to draw representative neurons.

The number of Prox1+ and Ki67+ cells in dentate gyrii were quantified by creating a pipeline in CellProfiler (http://cellprofiler.org). Images were first masked such that only the dentate gyrus was visible, then split into individual files by RGB channel (DAPI-blue, ki-67 – green, Prox1-red). Primary objects were identified for each channel using the following parameters: For nuclei, min-diameter 8, max-diameter 14, threshold correction 1.5, distinguishing by shape. For ki67, min-diameter 5, max-diameter 25, threshold correction 1, distinguishing by intensity. Prox1 primary objects were identified using the same settings as nuclei. Primary objects were then related to nuclei, removing false signal, and count was exported to a csv file. To calculate cells/μm, the area of the dentate gyrus measured was quantified manually in Fiji/ImageJ ^79^ (https://imagej.nih.gov/ij/).

The percentage of area occupied by CD68 inside Iba1+ microglia was quantified by creating a pipeline in CellProfiler and analyzing images in three batches. Images were first corrected for illumination, then aligned, and then Iba1 primary objects were identified with min pixel size 4 max 150 using an adaptive Otsu threshold strategy with three classes, identifying the foreground and a 0.1 lower threshold bound and an adaptive window size of 65. Second and third batches of CD68 stains required minor adjustments to the lower threshold bound for background correction.

### Behavioral Experiments

All behavioural experiments were undertaken in the Animal Outcome Core Facility (AOCF) at the Charité. Behavioral tests were performed within the guidelines granted by the LaGeSo under an extension to the experimental license G0079/11 and G0206/16. Prior to behavioral testing, male HP1FEC mice were implanted with subdermal RFID transponders to ensure accurate identification. Behavioral experiments were performed on both young adults (3 months) and aged adults (13-14 months). Prior to each cohort of behavioral testing, all animals were subjected to a modified **S**mithKline Beecham, **H**arwell, **I**mperial College, **R**oyal London Hospital, **p**henotype **a**ssessment (SHIRPA), which ensured animals did not have any gross deficits in vision, audition, grip strength, pina reflex and normal exploratory locomotion. After SHIRPA assessment, behavioral tests were always carried out at the same time of day, with tests spanning a 1 month period. Tests always occurred in the following order: Open Field Activity, Paired-Pulse inhibition, Context-Cued Fear Conditioning, Barnes Maze, Social Activity Monitor, HomeCageScan, Nest Construction.

#### Open Field Activity

Animals were placed in the centre of a square enclosure for 10 minutes while an overhead camera records and movement is tracked using the Biobserve Viewer Software. Activity in the ‘center zone’ and periphery were binned per minute.

#### Paired-Pulse Inhibition

Animals were tested two at a time in a 2-box startle box apparatus (TSE systems), which consisted of black soundproofed plexiglass boxes (49cm x 49cm x 49cm). The floors of internal cases were composed of metal bars connected to pressure sensors, which enabled precise measurement of startle response. Upon program start, animals acclimatized for 5 minutes, followed by a program (randomized by trial) that tested response to startle pulse alone (120dB for 40ms) or response to the pulse that had been preceded by a pre-pulse (one of 69dB, 73dB or 81dB for 20ms).

#### Context-Cued Fear Conditioning

Animals were analysed two at a time in two adjacent multi-conditioning boxes from TSE Systems. These were designed to be sound proofed and have a sound probe, a camera, and a context arena with a metal grated floor and walls high enough to contain the animal. Context cued fear learning was tested over 5 phases: Phase 1 (Shock): Animals were allowed to acclimatize in the chamber for 4 minutes (Context), which is followed by a 30 second sound (Cue), which is immediately followed by a weak electric shock on the floor of the context arena. Phase 2 (Context 24h): The following day, animals were placed back into the context arena and their freezing response was recorded over 3 minutes. Phase 3 (Cue 24h): two hours following phase 2, a cover was placed over the metal bar floor, animals were habituated for 3 minutes and then the cue sound was played for 3 minutes and freezing response was recorded. Phase 4 (Context 7 Days): Exactly as in phase 2, context was tested 7 days following phase 1. Phase 5 (Cue 7 Days): Two hours following Phase 4, animals were tested for freezing response to the cue, as in phase 3.

#### Barnes Maze

The Barnes Maze is a circular platform containing 20 holes around the circumference, one hole contains a submerged nest that serves as an escape from the open environment. Animals were trained on the location of the submerged nest. This involved placing the animal in the centre of the platform while loud static noise is played over four 3-minute trials over 4 training days, in which time the mouse could often find the submerged nest. If the mouse could not find the nest by the end of the 3 minutes, the mouse was shown the nest. Once in the nest the mouse was allowed to stay for 30 seconds to allow for positive reinforcement. Following 4 days of training, the nest is removed and animals were tested on the 5^th^ day (24h test) where mice were placed in the centre of the platform and their hole seeking behavior is recorded (time to target hole, errors before target hole, headpokes per hole) over a 90 second testing period. Animals were then tested one week later (7day test) in the same manner.

#### Social Activity Monitor (SAM)

To measure basic interaction and circadian activity, animals in their home cages were placed on top of RF sensors that detect motion. Animal activity was then recorded for 14 days using Phenoscoft control software. SAM data was binned by hour and following data export, RFIDs were decoded to corresponding animal ids and genotypes.

#### HomeCageScan (Microbehaviors)

Animals were recorded individually over 24hs using CleverSys Software for any changes in sterotyped murine behavior. Prior to recording, background cage registration, night/day and transition calibrations were set according to each cage. For data export, data was binned by minute and by hour.

#### Nest Construction

Because a pilot experiment revealed that a two day separation of male animals from their homecages resulted in hyper aggression upon their return, nest construction was only tested in the aged timepoint. To test nest construction ability, mice were housed individually and given a square piece of densely woven cotton called a ‘nestlet’ (Ancare). Animals were allowed to habituate with the nestlet for the first 24 hrs. For the second 24hs a new nestlet was supplied and the following morning what remained of the nestlet was weighed and the complexity of the nest was scored based on the standard rubric: (1) The Nestlet is largely untouched (>90% intact). (2) The Nestlet is partially torn up (50-90% remaining intact). (3) The Nestlet is mostly shredded but often there is no identifiable nest site: < 50% of the Nestlet remains intact but < 90% is within a quarter of the cage floor area, i.e. the cotton is not gathered into a nest but spread around the cage. Note: the material may sometimes be in a broadly defined nest area but the critical definition is that 50-90% has been shredded. (4) An identifiable, but flat nest: > 90% of the Nestlet is torn up, the material is gathered into a nest within a quarter of the cage floor area, but the nest is flat, with walls higher than mouse body height (curled up on its side) on less than 50% of its circumference. (5) A (near) perfect nest: > 90% of the Nestlet is torn up, the nest is a crater, with walls higher than mouse body height on more than 50% of its circumference.

### Magnetic Resonance Imaging

MRI was performed at a 7 Tesla rodent scanner (Pharmascan 70 ⁄ 16, Bruker, Ettlingen, Germany) with a 20 mm diameter transmit/receive volume resonator (RAPID Biomedical, Rimpar, Germany). For imaging the mouse brain a T2-weighted 2D turbo spin-echo sequence was used (imaging parameters TR/TE = 5505 ms/36 ms, rare factor 8, 6 averages, 46 axial slices with a slice thickness of 0.350 mm, field of view of 2.56 x 2.56 cm, matrix size 256 x 256; scan time 13m12s). MRI data were registered on the Allen mouse brain atlas (ABA) using an in-house developed MATLAB toolbox ANTx (latest version available under https://github.com/ChariteExpMri/antx2). The volumes of each single ABA brain structure were calculated using the back-transformed atlas which matched the individual T2-weighted images ^80^. For section-wise analysis, the mean isocortex volume per section per genotype per age was calculated and tested using two-way ANOVA.

### Site specific 5hmC and 5mC Analysis by qPCR

Genomic DNA was purified from frozen hippocampi using NucleoSpin Tissue columns (Macherey-Nagel) according to the manufacturer’s instructions. Purified gDNA was subsequently processed using the EpiMark 5hmC Analysis kit (NEB, cat# E3317) according to the manufacturer’s instructions. CpGs of enzymatically prepared samples were then profiled using primers designed to amplify over an area containing a single HpaII/MspI site. Primers used for qPCR: Nnat: ACCCCTCCTTCTCAACATCC & CGCCGAGGTCTACTGGTCT. For IAPEz: CTTTGAAGGAGCCGAGGGTG & AAGCCTGTCTAACTGCACCAA. qPCR was performed using the GoTaq qPCR Master mix (Promega) including the CXR reference dye on a StepOnePlus Thermocycler (Applied Biosystems).

### HP1cTKO Embryonic Stem Cell line

ESCs possessing *Cbx1*, *Cbx3*, or *Cbx5* conditional alleles were constructed via gene targeting by flanking exons 2 and 3, exon 3, or exon 3, of each gene with LoxP sequences, respectively (fig. S12). Conditional mice were established from each conditional ESC line. *Cbx1*, *Cbx3*, or *Cbx5* mutant mice were next crossed with mice bearing *CreERT2* alleles, to enable excision of the floxed alleles by addition of 4-OH tamoxifen (4-OHT). Triple conditional mutant mice were obtained via crossing the single, or double conditional mice. Triple conditional ESC lines were made in house from the triple conditional mouse blastocysts. ESCs were cultured in D-MEM (Kohjin-bio, #16003550) with 20% fetal bovine serum (Sigma-Aldrich, #172012), MEM nonessential amino acids (GIBCO, #11140-050), L-glutamine (GIBCO, #25030-081), 2-mercaptoethanol (Sigma-Aldrich, #M1753), and LIF (in-house preparation) on mitomycin C-treated (Sigma-Aldrich, #M4287) primary MEF feeder layers. For conditional KO, 4-OH Tamoxifen (SIGMA, #H7904; (Z)-4-Hydroxytamoxifen, ≥98% Z isomer, dissolved in ethanol) was added to medium to a final concentration of 800 nM and cultured for 6 days. The stock solution was prepared as 2 mM (X2500).

### Chromatin Immunoprecipitation (ChIP) & Library Construction

ChIP experiments and subsequent library preparation, were performed as previously described ^81^. Antibodies used for ChIP can be seen in Data S1.

### Reduced Representation Bisulfite Sequencing (RRBS)

RRBS was carried out following a previously described method ^82^ with minor modifications. 500 ng of genomic DNA was used as a starting material. Bisulfite conversion was done by the EZ DNA Methylation Gold Kit (Zymo Research, #D5005) with 50 ng of DNA, per sample. 2x KAPA HiFi Hot Start Uracil+ Ready Mix (KAPA Biosystems, #KK2801) was utilized for library amplification. PCR amplification was done for 10 cycles.

### 5mC & 5hmC Joint profiling from Hippocampal Lysates

Hippocampi were isolated in ice cold PBS and DNA was using the QIAamp Fast DNA Tissue Kit (QIAGEN, cat # 51404). Purified DNA was then used to generate bisulfite converted or oxidative bisulfite converted RRBS libraries using the Ovation RRBS kit (TECAN, formerly NuGEN, cat # 0553-32). RRBS libraries were sequenced in a 50bp paired end configuration with an additional 6bp allotted to the library index on an Illumina Novoseq 6000.

### Bioinformatics & Data Processing

Bioinformatic pipelines were written using Snakemake https://snakemake.readthedocs.io/en/stable/ and deployed on the cluster hosted by the Berlin Institute of Health (BIH). Scripts for analysis are provided at https://github.com/qoldt/HP1-Deficiency-Neurodegeneration.

For RNAseq data, fastq files were aligned to GRCm38.p5 using STAR ^83^ with the following settings to maximize repeat mapping (--outFilterMultimapNmax 100, --winAchnorMultimapNmax 100, --outSAMstrandField intronMotif). The TETranscripts ^84^ package was used to generate a count table using gencode.vM16.basic.annotation.gtf and the prepared repeat masker file. The outputted count table was re-annotated using biomaRt ^85^ and analyzed for differential expression using edgeR ^86^. Hierarchical clustering and heatmap of significantly changed transcripts determined from testing between all WT and all HP1DKO (689 genes, adjusted p < 0.05, Data S1) was created from a scaled matrix using heatmap2.

For IAP RNAseq coverage profiles, known IAP coordinates were obtained from the UCSC table browser in bed format. Deeptools ^87^ was used to generate RPKM normalized 50bp bin bigwig files with the option --extendReads from aligned RNAseq data. Comatrices were computed to scale regions to an internal size of 500bp with before and after region lengths of 1000bp. Read coverage was plotted over IAP elements for each genotype at each age using RPKM (per bin) = number of reads per bin / (number of mapped reads (in millions)* bin length (kb)). Chimeric transcripts were detected using a combination of LIONS ^88^ and use of a 1000bp running window filter in Seqmonk. Inflammatory response was profiled by cross referencing genes differentially expressed in HP1DKO with the interferome database ^89^. Gene set enrichment and leading edge analysis was performed using GSEA to query raw count data from aged HP1DKO and wildtype against c2.cp.reactome.v6.2.symbols.gmt using Signal2noise in gene_set mode with default parameters. GSEA output was imported into cytoscape using the EnrichmentMap ^90^ plugin (Jaccard Overlap combined cut-off = 0.375, k constant = 0.5, node cut-off Q = 0.6 and edge cutoff similarity of 0.53). Network node clusters were coarsely annotated using the AutoAnnotate ^91^ plugin which was further refined using Adobe Illustrator.

5mC and 5hmC RRBS data derived from paired bisulfite (BS) and oxidative bisulfite (oxBS) reactions was analysed as follows: Reads (R1 and R3 in this case) were trimmed using trim_galore (https://github.com/FelixKrueger/TrimGalore) with the parameters –-paired -a AGATCGGAAGAGC -a2 AAATCAAAAAAAC. Reads were then further processed using the NuGEN diversity trimming script (https://github.com/nugentechnologies/NuMetRRBS/blob/master/trimRRBSdiversityAdaptCustomers.py) and aligned to the GRCm38 (mm10) Bisulfite Genome using Bismark with -p 2 -N 1 --multicore 8. Coverage files were generated using bismark_methylation_extractor with -p --ignore_r2 3. True 5mC was taken directly from the oxBS data. To infer 5hmC state, oxBS coverage files were subtracted from BS coverage files outlined in the created ‘Extract 5hmC.R’ script, where count *Count_5hmC_* = *Count_BSmethylated_* − *Count_oxBSmethylated_*, %*_5hmC_* = %*_BSmethylated_* − %*_oxBSmethylated_*, and the number of unmodified cytosines *Count_No5hmC_* is calculated based on the relationship 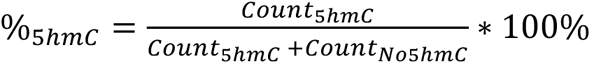, rounded to the closest integer. In fringe cases where oxBS signal is higher than BS (5hmC is negative), 5hmC is set to zero and *Count_No5hmC_* is set to *Count_BSunmethylated_*. Coverage files from 5mC and 5hmC were tested for differential methylation using methylkit ^92^ and intersected with genomic annotations obtained from the UCSC table browser. Odds ratio for each annotation tested was calculated as follows; given *D_overlaps_*, the number of Differentially Methylated Regions (DMRs) that overlap with the annotation, *N*_overlaps_, the number of Non-Differentially Methylated Regions (Non-DMRs) that overlap with the annotation, *D*, the total number of DMRs and *C*, the total number of observed cytosines:

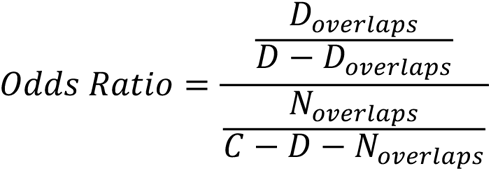

The probability *P* of drawing *D_overlaps_* or more by chance, when drawing *D* DMRs from a population of *C* total cytosines, of which *D_overlaps_* and *N*_overlaps_ are successes (i.e., overlaping with the annotation) was calculated using the hypergeometric distribution. This calculation was performed with the ‘phyper’ function in R which tests the null hypothesis that the observed number of overlaps is as expected by chance: *P*(*X* ≥ *D*_*overlaps*_) = 1 − *phyper*(*D*_*overlaps*_ − 1, *D*_*overlaps*_ + *N*_*overlaps*_, *C* − (*D*_*overlaps*_ + *N*_*overlaps*_), *D*, *lower.tail* = *TRUE*). Following the hypergeometric test, p values were adjusted for multiple comparisons using the Benjamini-Hochberg procedure based on the number of annotations tested per dataset. Summary plots of methylation were generated using the plotAnnopeak function of ChIPseeker and Heatmaps of DMRs were generated using ComplexHeatmap^93^.

HP1cTKO RRBS data was quality trimmed using trim galore using the --rrbs flag and prepared for analysis using Bismark ^94^ with bowtie1 and arguments -n 1 -l 45. Bismark coverage files from WT and HP1cTKO were analysed using the R package methylKit ^92^, where all differentially methylated bases were extracted that change more than 25% and pass a corrected significance threshold of q = 0.01. Differentially methylated bases were then annotated using genomation ^95^ and prepared bed files (CpGs, LINEs, SINEs, LTRs, Exons) retrieved from the UCSC table browser. Jitter plots were created using ggplot2. Circos plot was prepared by using the circlize package ^96^. HP1cTKO eAge was calculated by using the ∼18,000 CpGs included in the MouseEpigeneticClock tool (https://github.com/EpigenomeClock/MouseEpigeneticClock) ^97^. In the WT sample 966 (5.36%) sites were imputed whereas in HP1cTKO 946 (5.25%) sites were imputed. The original eAge prediction for WT was -0.9 weeks and the HP1cTKO 0 weeks resulting in a positive difference of 0.9 weeks (6.3 days).

ChIP raw data was quality filtered using Trimmomatic and aligned to GRCm38.p5 using bowtie. Deeptools was used to generate bigwig files and generate heatmaps and profiles ^87^. RPGC normalized bigwig files were created per condition by averaging biological replicates using bamCompare with parameters --normalizeUsing RPGC --effectveGenomeSize 2652783500 -- operation mean --extendReads 125. Comatrices were computed to scale *IAPEz-int* regions to 1kb in addition to taking 1kb upstream and 1kb downstream. KAP1, H3K9me3 and H4K20me3 binding profiles were quantified by creating a centre-point comatrix at ZFP57 peaks overlapping published murine ICRs in deeptools. For this annotation, published ZFP57 peaks ^42^ were subset by their overlap with published murine ICRs.

## Supplementary Text

### Behavior

For behavioral experiments, A total of 42 animals (15 WT, 11 HP1βKO, 6 HP1γKO, 10 HP1β/γDKO) completed the aged time point, while 4 died (2 WT, 1 HP1βKO, 1 HP1β/γDKO) between young and aged testing. Box and whisker plots for behavioral experiments are comprised of median and 25^th^ and 75^th^ percentiles, where whiskers extend no further than 1.5X the interquartile range. All line charts plot the mean, with standard error rendered as a ribbon surrounding the line. Unless stated otherwise, behavioral experiments were analyzed using Two way ANOVA with Bonferroni correction for multiple comparisons; where Asterisks (*) denote tests to between genotype within age (* = p < 0.05, ** = p < 0.01, *** = p < 0.001) and hashtags (#) denote tests within genotype between age (# = p < 0.05, ## = p < 0.01, ### = p < 0.001).

### HP1cTKO Cell line

After 7 days of tamoxifen (4-OHT) induced deletion of HP1 proteins in the HP1cTKO cell line, the cells failed to thrive and cell division became extremely slow. We suspect that changes to 5hmC and 5mC in HP1cTKO ESCs at 6 days are in an intermediate stage, similar to that seen previously in SETDB1 -/- ESCs which were cultured for 4 days ^98^.

### RNAseq Longitudinal Analysis

No Differentially expressed genes could be detected between young HP1γKO and aged HP1γKO. Young HP1βKO and aged HP1βKO showed 184 differentially expressed transcripts among which were increases in C4 and C1qa in aged HP1βKO. A non-negligible batch effect meant direct comparison of young HP1DKO and aged HP1DKO was not statistically advisable.

---

**Figure S1.**
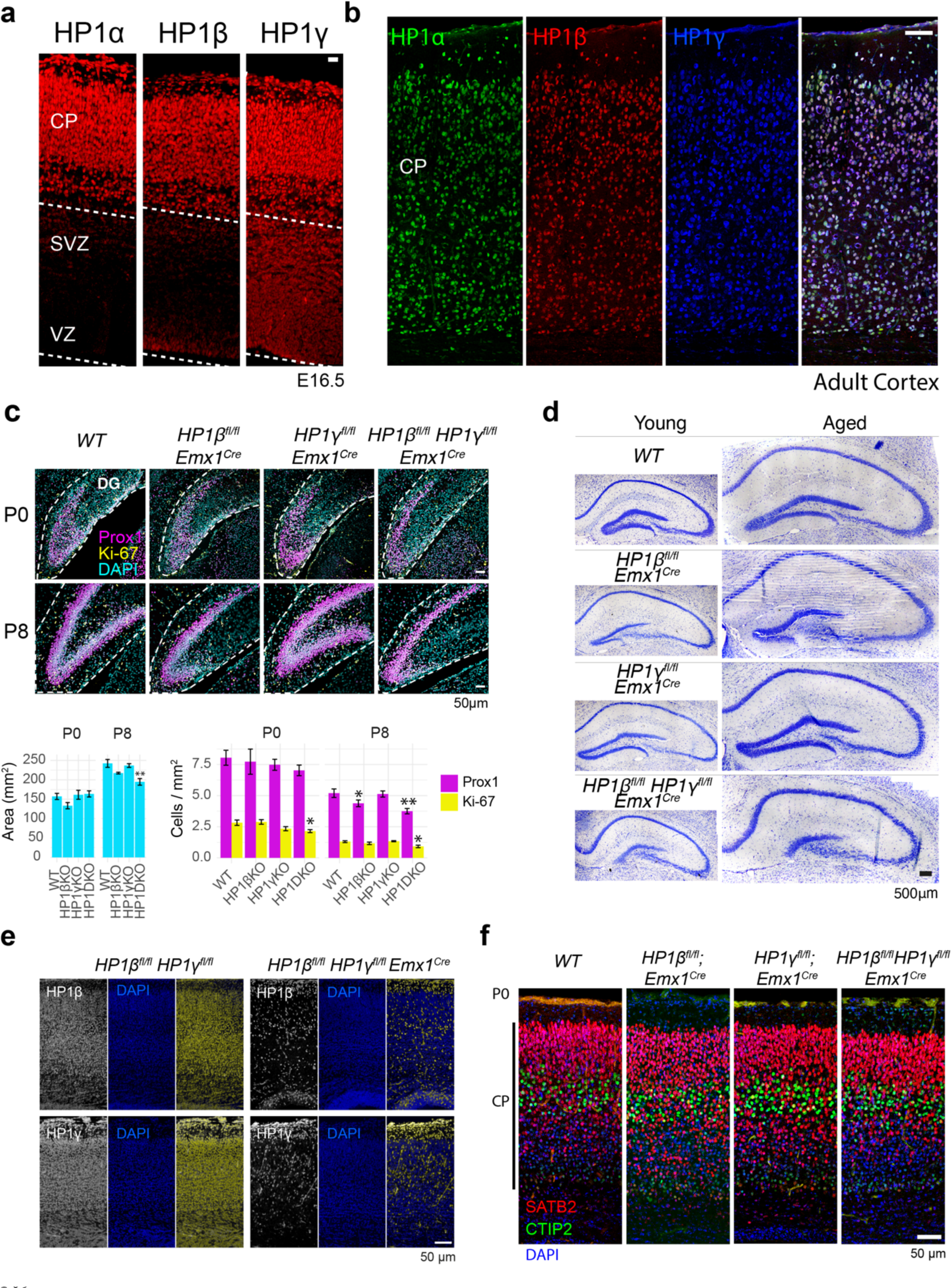
Expression of HP1 proteins and deletion from the cerebral cortex. **a** HP1α and HP1β expression is strongest in post-mitotic neurons of the cortical plate (CP) at embryonic day 16.5. Both are also expressed in progenitor cells in the Subventricular and ventricular zones (SVZ and VZ, respectively). HP1γ expression is much more uniformly expressed across progenitor and post-mitotic cell types. (scale bar = 50 μm) **b** HP1 proteins remain robustly expressed in the cortical plate of adult mice (scale bar = 100μm). **c** Dentate gyri of HP1βKO and HP1β/γKO animals show malformation of the infrapyramidal blade due to mitotic exhaustion between postnatal day 0 and postnatal day 8. P0 WT n = 10, P0 HP1βKO n = 4, P0 HP1γKO n = 7, P0 HP1DKO n= 11, P8 WT n = 4, P8 HP1βKO n = 3, P8 HP1γKO n = 4, P8 HP1DKO n= 5. One way ANOVA with Dunnett’s Multiple Comparison test where ** denotes p < 0.05, and * denotes p < 0.01. **d** The morphology of young (3 month) and aged (13 month) hippocampi. **e** Emx1-Cre mediated deletion of floxed HP1 proteins results in targeted deletion of HP1β and HP1γ in the pyramidal lineage seen here at postnatal day 0 (P0). **f** Single or double deletion of HP1β or HP1γ does not adversely affect neuronal cell fate or laminal position in the cerebral cortex at P0.

**Figure S2.**
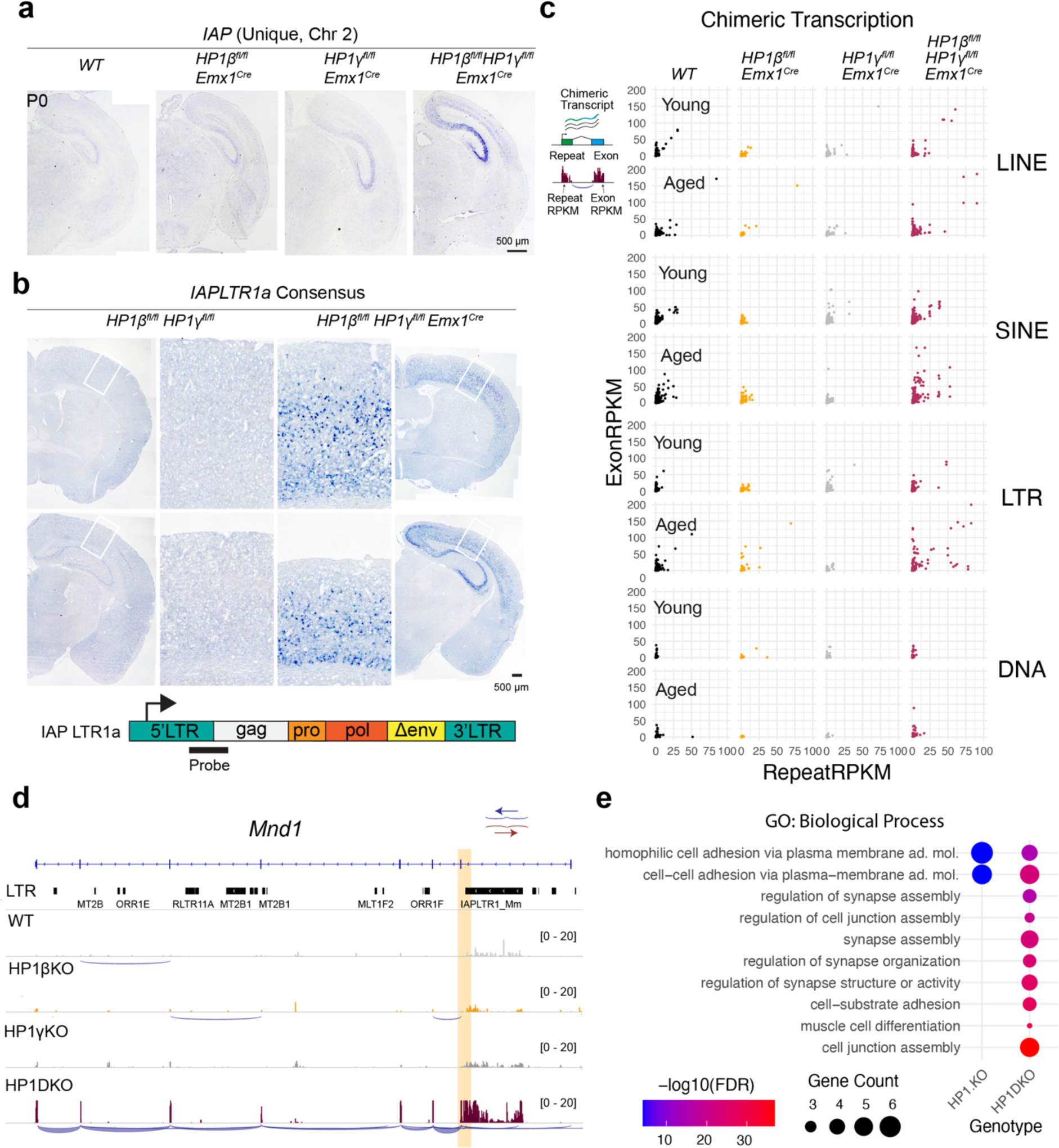
Sustained de-repression of ERVs and induction of chimeric transcripts in HP1DKO mutants. **a** In situ hybridization on P0 brains using an RNA probe specific for a single IAP element on chromosome 2. **b** In situ hybridization on adult brains using an RNA probe that recognizes an IAPLTR1a consensus sequence found on at least 196 IAPs. **c** Co-transcription plots of chimeric transcripts where each point represents a chimeric transcript. **d** An example chimeric transcript; transcriptional activation of an IAP element in the first intron of *Mnd1* results in transcriptional activation of the entire gene. Shown here are RNAseq tracks from aged genotypes. **e** Gene Ontology of significantly changed genes (edgeR FDR <0.05) in *HP1β^fl/fl^Emx1^Cre^*, *HP1γ^fl/fl^ Emx1^Cre^*, and *HP1β^fl/fl^ HP1γ^fl/fl^ Emx1^Cre^* RNAseq. No significant gene ontology could be observed for *HP1β^fl/fl^Emx1^Cre^*. The differential expression of protocaherins in *HP1γ^fl/fl^ Emx1^Cre^*, and *HP1β^fl/fl^ HP1γ^fl/fl^* greatly biases gene ontology towards cell adhesion and synapse gene sets.

**Figure S3.**
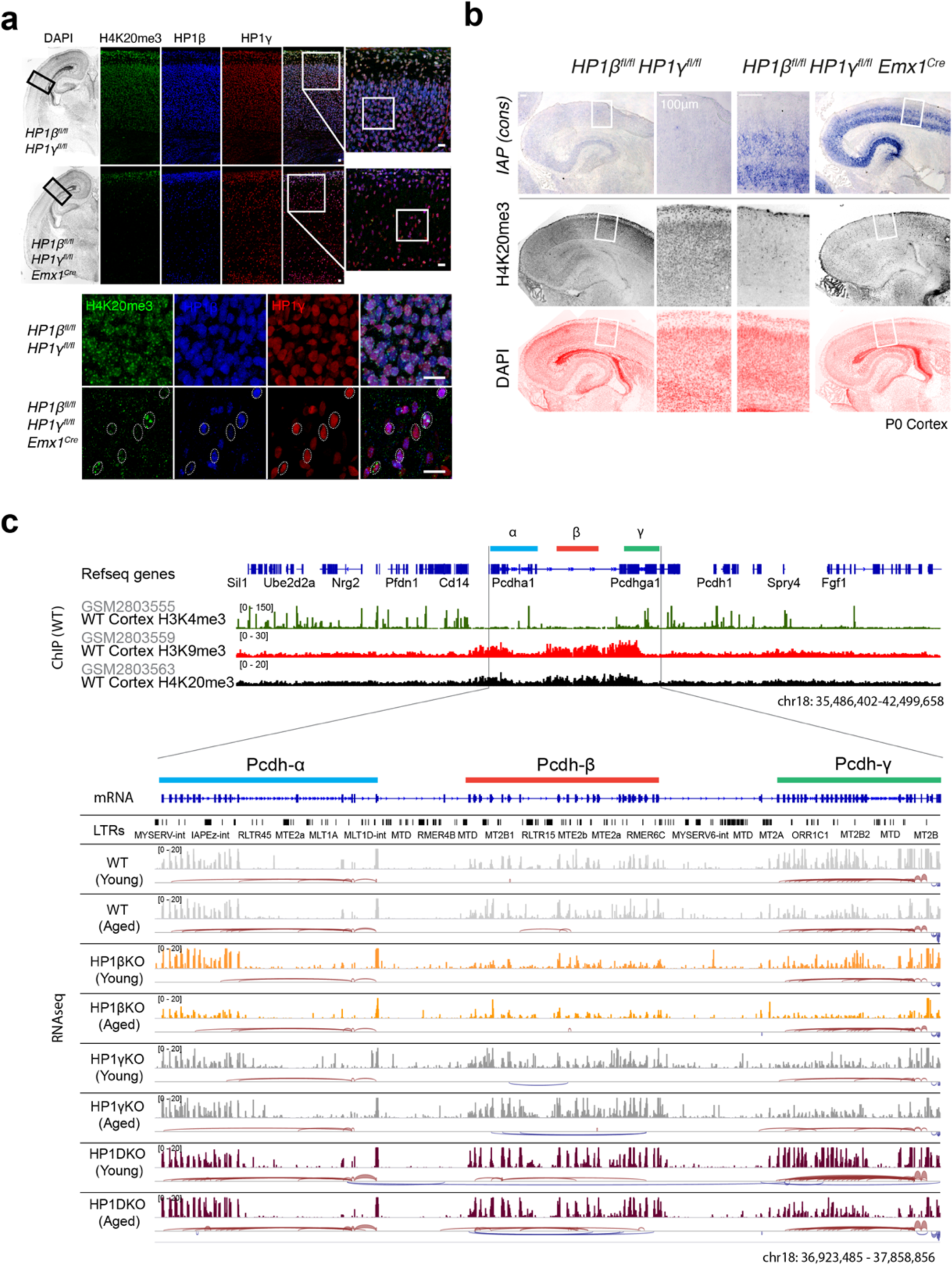
H4K20me3 loss is correlated with IAP de-repression and Protocadherin expression. **a** H4K20me3, normally abundant in post-mitotic cortical neurons is completely missing in HP1γ deficient Emx1-lineage pyramidal neurons. In cortices missing HP1γ, H4K20me3 can still be seen in adjacent wild type interneurons (see magnifications). All scalebars 50μm. **b** *In situ hybridization* of IAP consensus sequence alongside adjacent sections stained for H4K20me3 and Dapi. **c** HP1γ is required for transcriptional regulation of the protocadherin cluster. (Top): Reference gene annotation of the mouse protocadherin α β and γ clusters on chromosome 18 along with published ChIPseq data of H3K4me3, H3K9me3 and H4K20me3 performed in cortical neurons ^41^. (Bottom): mRNA and LTR reference annotation along with read coverage tracks from hippocampal RNAseq performed in this study.

**Fig. S4.**
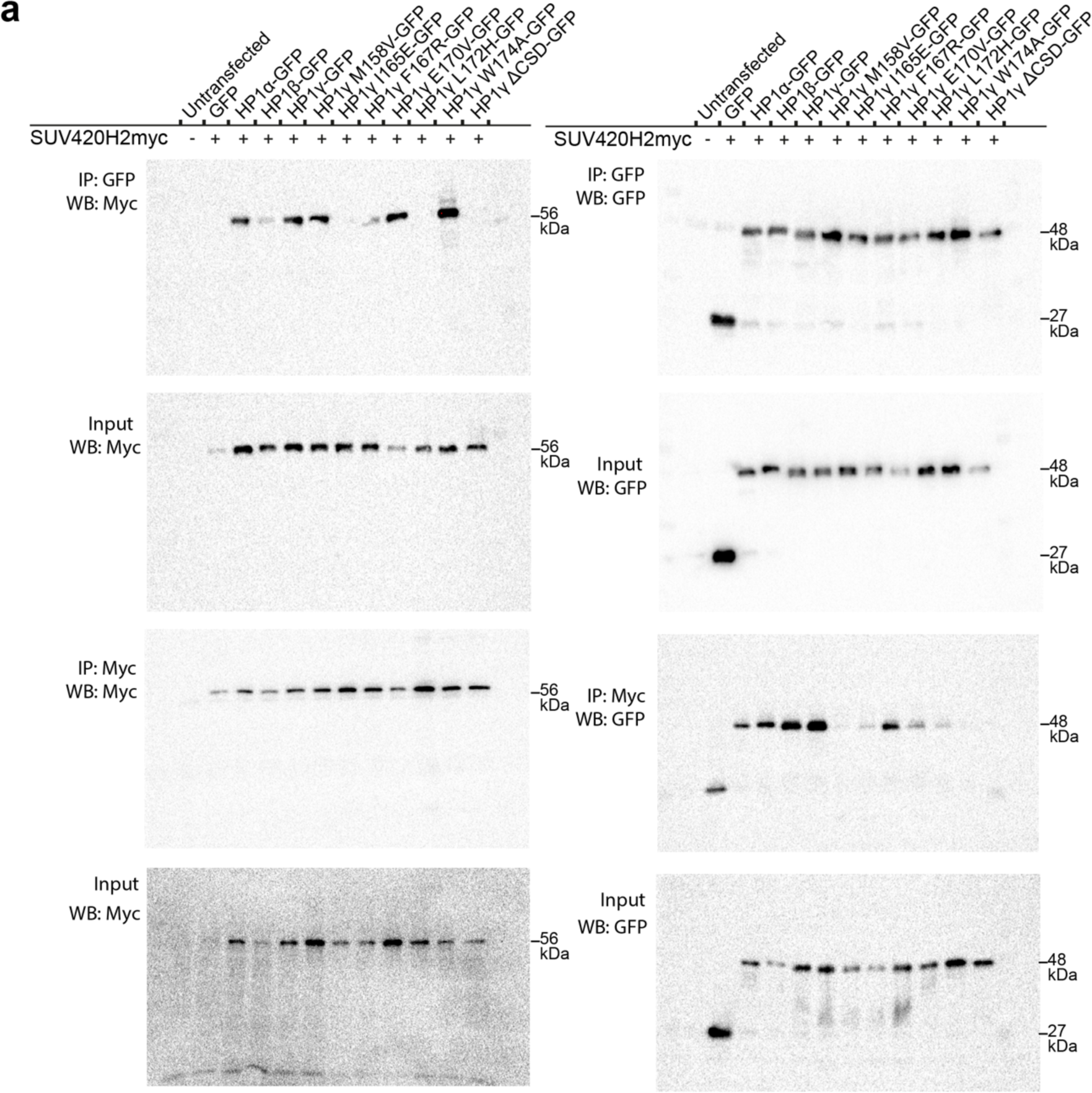
(B) Full blots from Co-immunoprecipitation of SUV420H2myc with HP1-GFP proteins from Figure **2d**.

**Figure S5.**
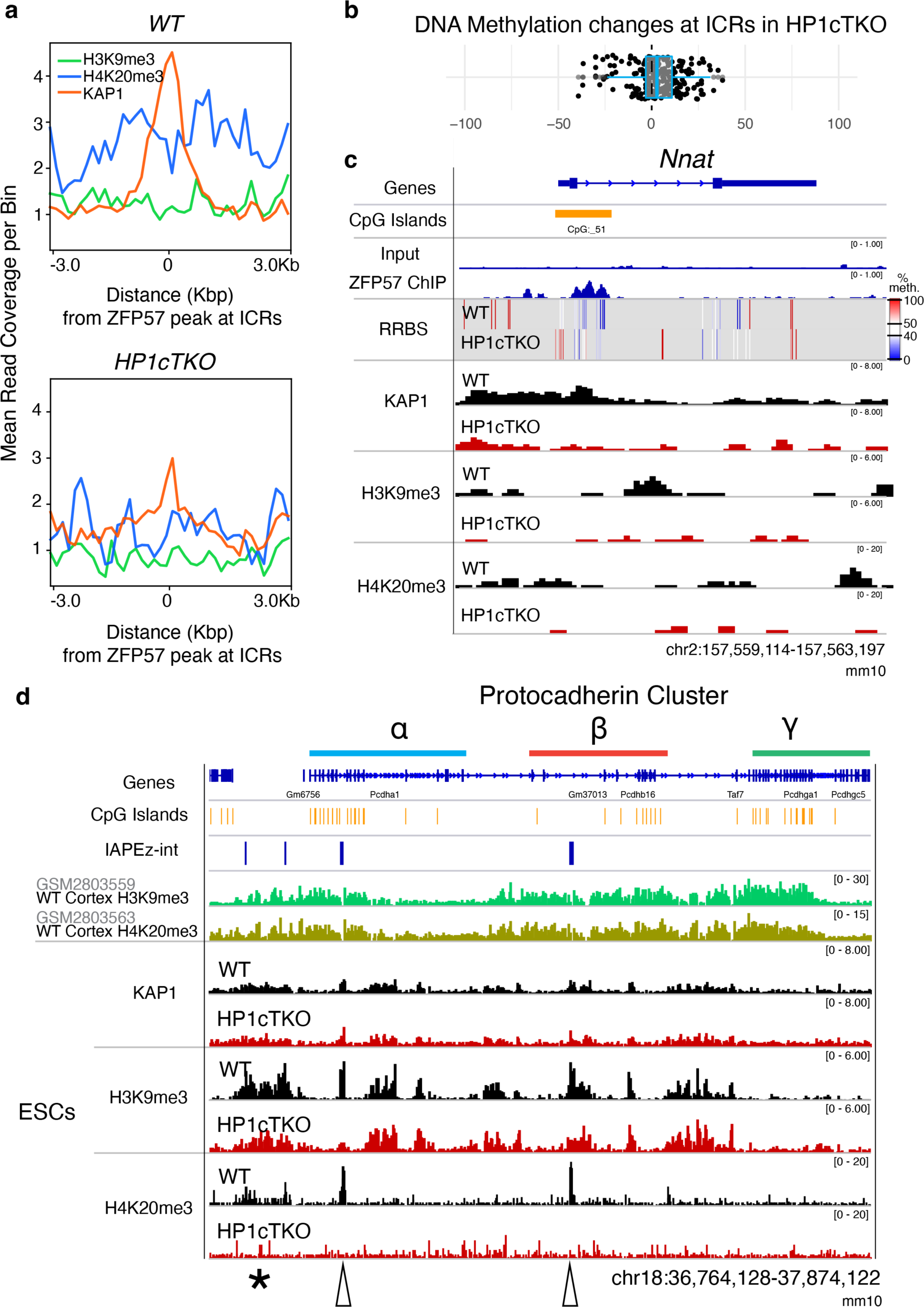
H3K9me3 is lost at repeats and ICRs but is unchanged at regulatory regions of the protocadherin cluster in HP1cTKO ES cells. **a** Read coverage of KAP1, H3K9me3 and H4K20me3 centred at ZFP57 peaks found at mouse imprinting control regions (ICRs) averaged over biological replicates. (RPGC-normalized bigwigs were averaged, 50bp comatrices were used and profiles were plotted over ICRs with a window of 150bp). **b** DNA Methylation changes at ICRs in HP1cTKO are mixed but show a tendency towards ‘hypermethylation’ which would be consistent with increased 5hmC observed in figure S5a. **c** Genome browser view of changes occurring in HP1cTKO ESCs at imprinting control region found at Neuronatin (*Nnat*). In addition to shifts in DNA methylation seen in the RRBS track, KAP1 is lost in HP1cTKO ESCs surrounding the ICR (CpG island marked by ZFP57; ZFP57 ChIP data from ^42^). **d** Genome browser view of changes to KAP1, H3K9me3 and H4K20me3 at the protocadherin locus in HP1cTKO ESCs with reference tracks for refseq genes, CpGs, IAPEz-int, along with published ChIPseq of H3K9me3 and H4K20me3 from the cortex ^41^. H3K9me3 and H4K20me3 mark regulatory domains over the protocadherin cluster in cortical neurons. At the same locus in embryonic stem cells (ESCs), H3K9me3 can be seen at IAPs and regulatory domains, and in HP1cTKO cells H3K9me3 is only lost at IAPs. H4K20me3 at the protocadherin cluster in wt ES cells is prominently observed at IAP loci (empty arrows) and at the proximal promoter (*), and is completely lost in HP1cTKO ESCs.

**Figure S6.**
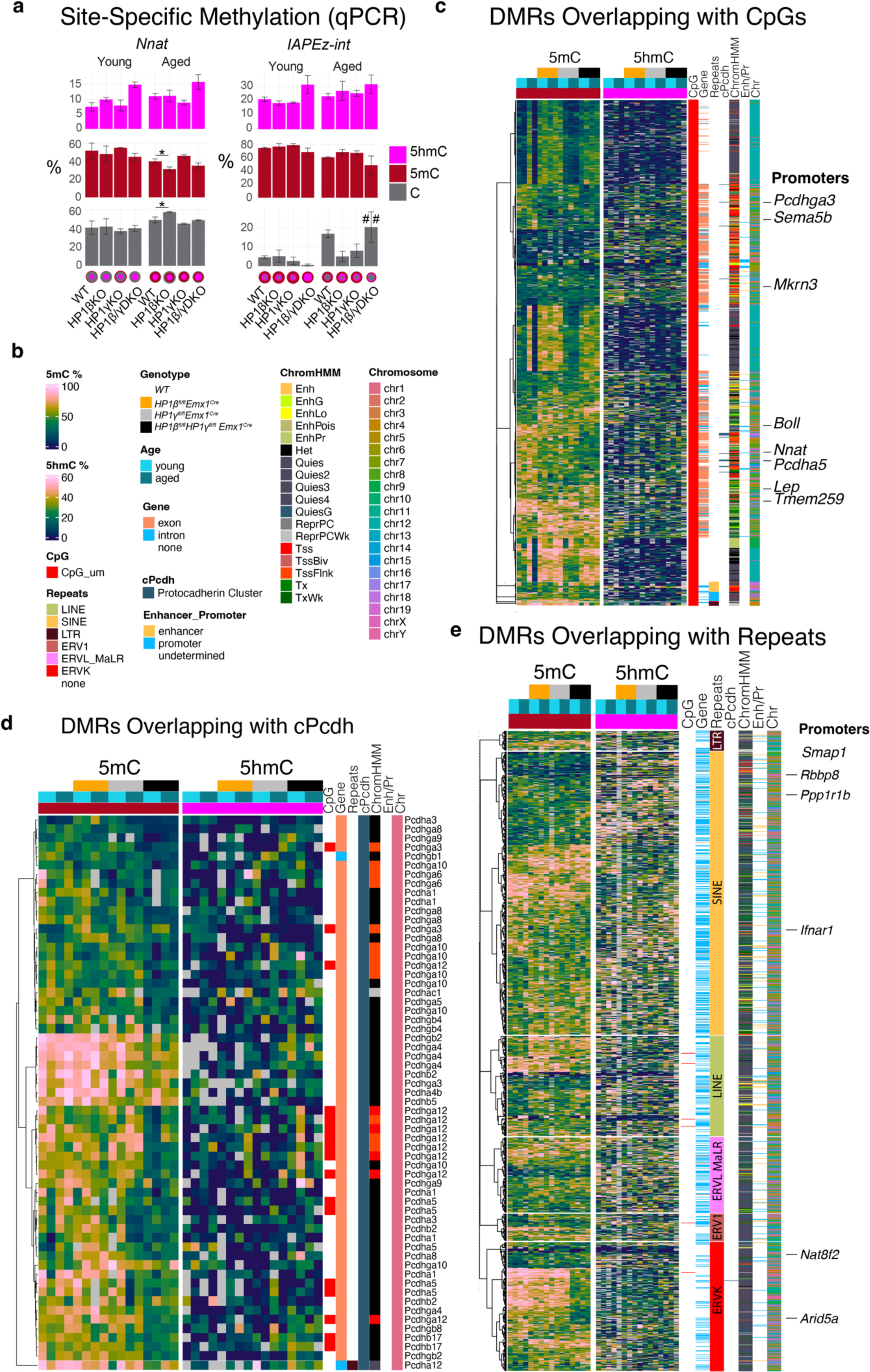
DNA methylation is affected in several regions in HP1 deficient hippocampi. **a** Profiles of qPCR-profiled cytosine hydroxymethylation (5hmC), cytosine methylation (5mC) and unmethylated cytosines (C) at selected CCGG sites in Neuronatin (*Nnat*) and IAPEz-int from hippocampal lysate DNA. Mean ±SEM; 3 biological replicates for each, two-way ANOVA, * = p <0.05 within age across genotype, ## = p <0.01 within genotype across age. **b** Legend for heatmaps in **c,d,e** for significant DMRs subset by row annotations corresponding to CpG islands, gene body, repeats, P0 Cortex ChromHMM state and chromosome. **c** DMRs (% difference > 25%, q < 0.05) overlapping annotated CpG islands. **d** DMRs (% difference > 25% q < 0.05) occurring in the protocadherin cluster. **e** DMRs (% difference > 30%, q < 0.05) that overlap an annotated repeat.

**Figure S7.**
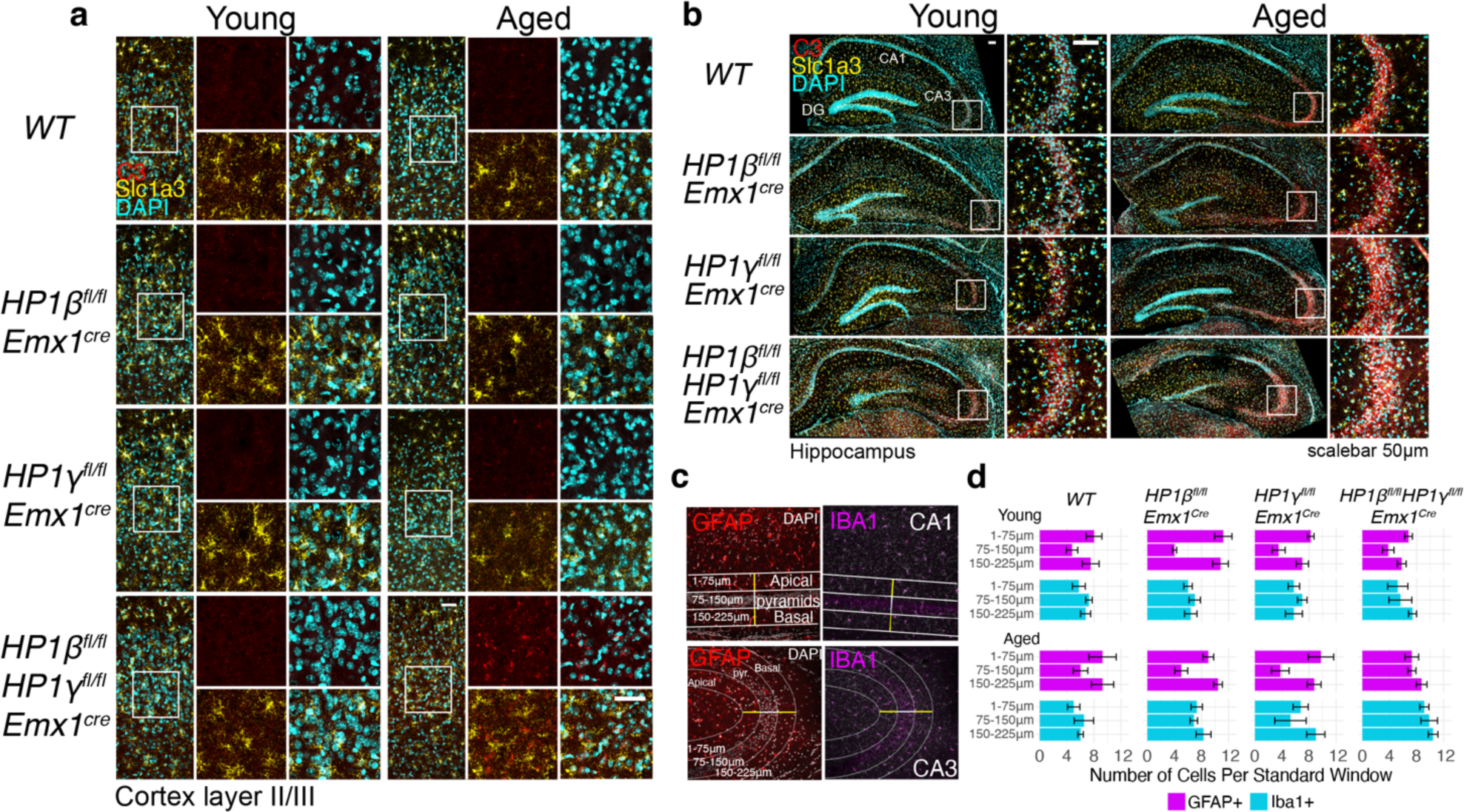
Initiation of Complement and invasion of microglia in HP1DKO cortices and hippocampi. Multiplex In situ hybridization using probes for *C3* and *Slc1a3* in the cortex (**a**) and hippocampus (**b**) of young and aged HP1 mutants. Distribution of astrocytes and microglia in or adjacent to pyramidal layers of the hippocampus. **c** Quantification scheme for CA1 and CA3 fields. Iba1+ microglia and GFAP+ cells with astrocyte morphology were quantified with 75 μm bins in standard windows across apical, pyramidal and basal fields. For CA1, straight lines separated by 75μm equivalent pixels was used, whereas for CA3, standard ellipses separated with 75μm equivalent pixels were used. Quantification summary of GFAP+ and Iba1+ can be seen by bin in **d** and across all fields in figure **5m**.

**Figure S8.**
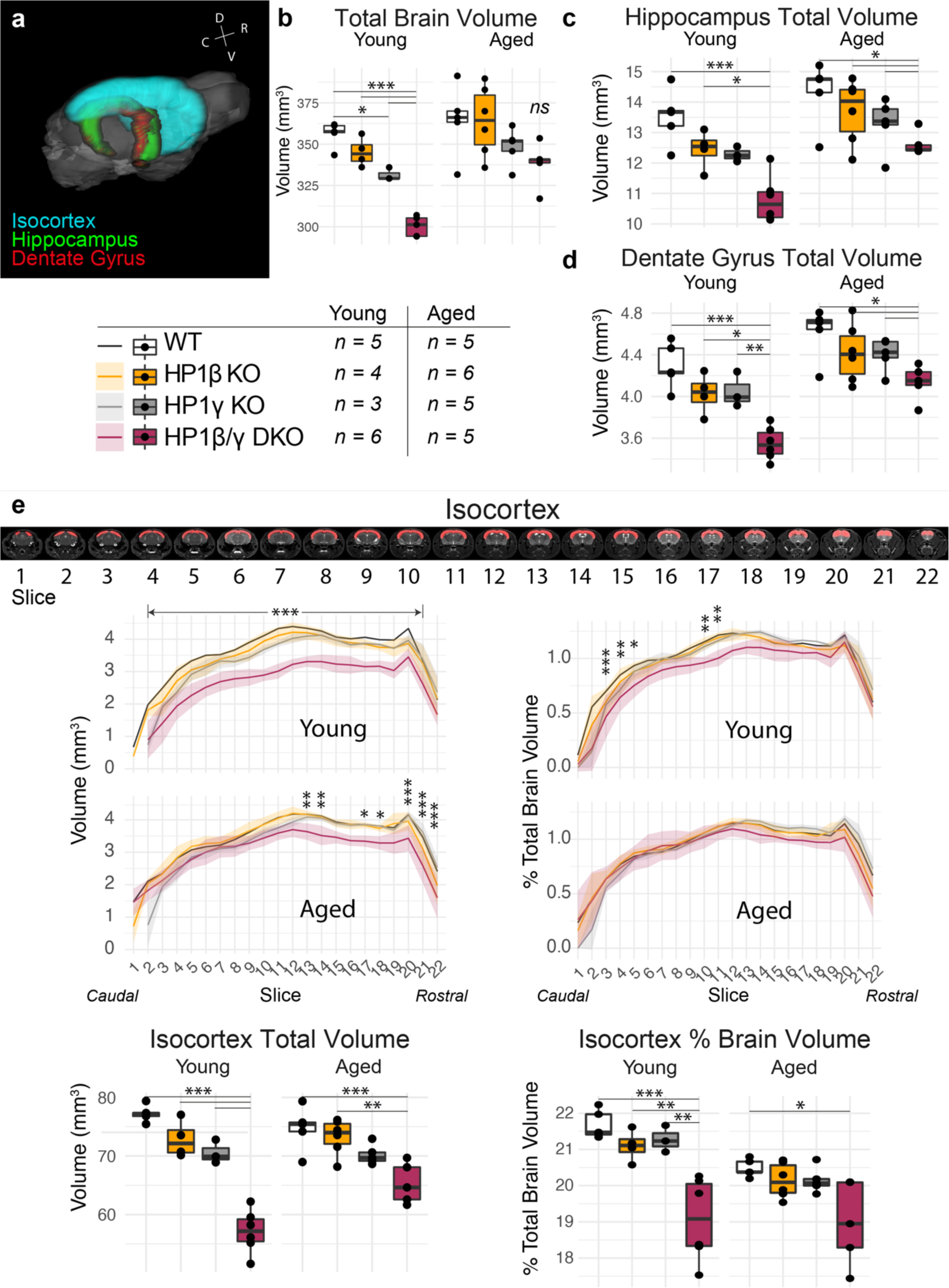
Structural Magnetic Resonance Imaging of HP1 mutants. **a** Registration of Isocortex, Hippocampus and Dentate Gyrus. Absolute volumes of total brain **b**, Hippocampus **c** and dentate gyrus **d** in young and aged HP1 mutants. **e** Slice-wise depiction of isocortical volume, both absolute (left) and as a percentage of total brain volume (right). All animals were included in the analysis. Statistics: one-way ANOVA within age with Bonferroni multiple comparison corrections. *** = p < 0.001, ** = p < 0.01, * = p < 0.05.

**Figure S9.**
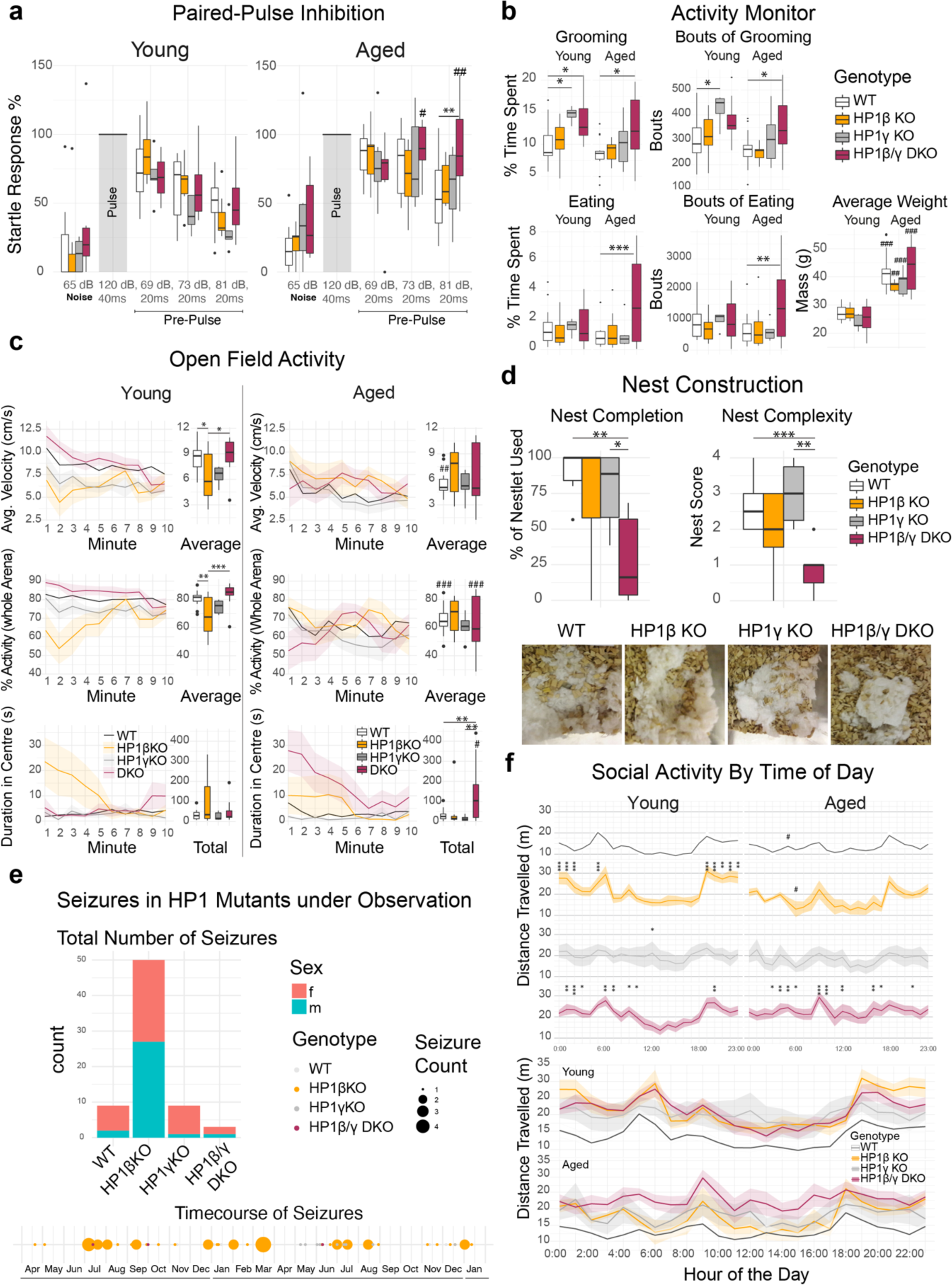
Age-related changes in behavior in HP1β/γDKO animals. **a** Paired-Pulse Inhibition recordings across HP1 mutants. All paired startle responses are percent normalized to the amplitude of the startle pulse (120dB) alone. A strong age-dependent change in paired pulse inhibition was seen in HP1β/γDKO animals, where pre-pulses of larger amplitude failed to further attenuate a startle response. **b** Altered Grooming, Eating and Circadian Activity in HP1mutants. HP1DKO animals showed an increased total time and frequency of grooming, and aged HP1DKO also displayed higher bouts of eating, though this did not translate into an increase in weight. Anxiety and exploration measured in an open field test revealed age dependent delays in HP1DKO animals (**c**). Young HP1βKO animals displayed non-convulsive freezing responses upon placement in the chamber which bias their measurements. **d** Nest construction profiled in aged animals was scored based on nest completion (% Nestlet used) and nest complexity (standard rubric, see methods) show marked deficits in HP1DKO animals. **e** Seizures observed in HP1 animals occurred predominantly in HP1β KO animals and were typically stimulation dependent, where observed seizures were non-randomly distributed, often occurring because of standard cage changes. Social activity **f** of mice observed over a 14 day period binned by hour reveals general hyperactivity in HP1βKO and HP1β/γDKO animals as well as an age-dependent change in peak nocturnal activity. Statistics are two-way ANOVA with Bonferroni correction on multiple comparisons; Asterisks (*) denote tests to wildtype within age, hashtags (#) denote tests within genotype between age.

**Extended Data Figure S10.**
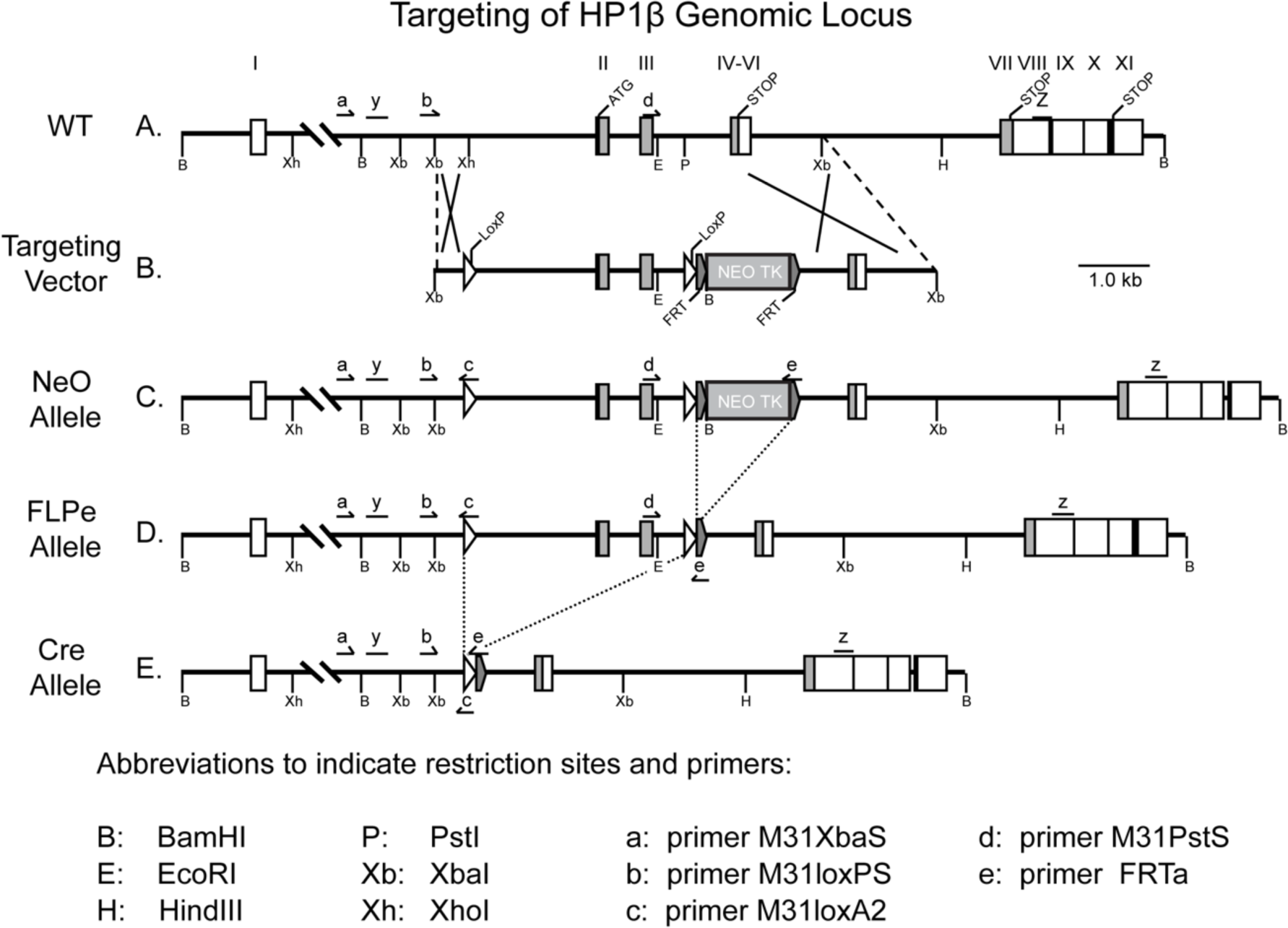
Targeting of Cbx1 (HP1β) genomic locus.

**Extended Data Figure S11.**
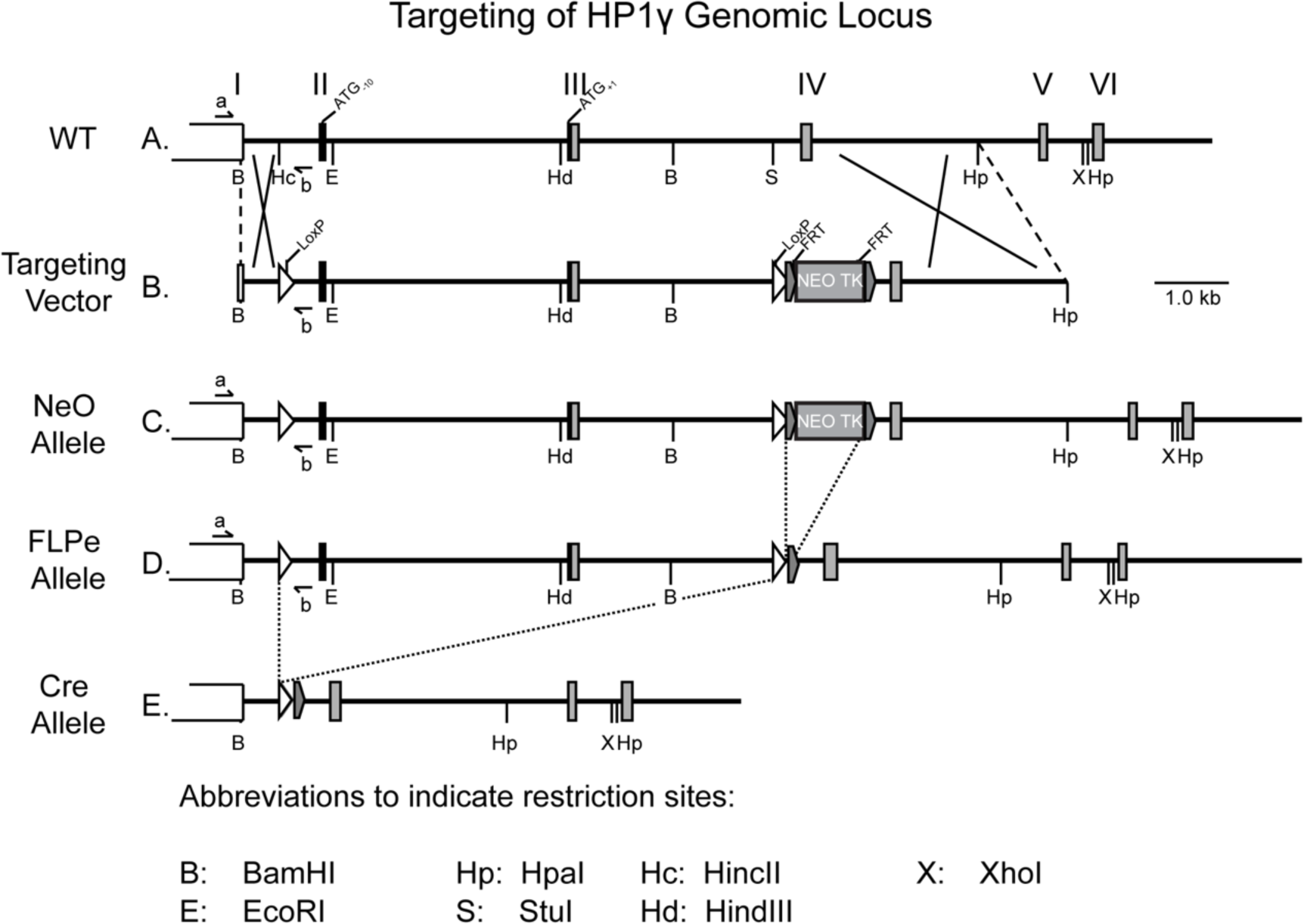
Targeting of Cbx3 (HP1γ) genomic locus.

**Extended Data Figure S12.**
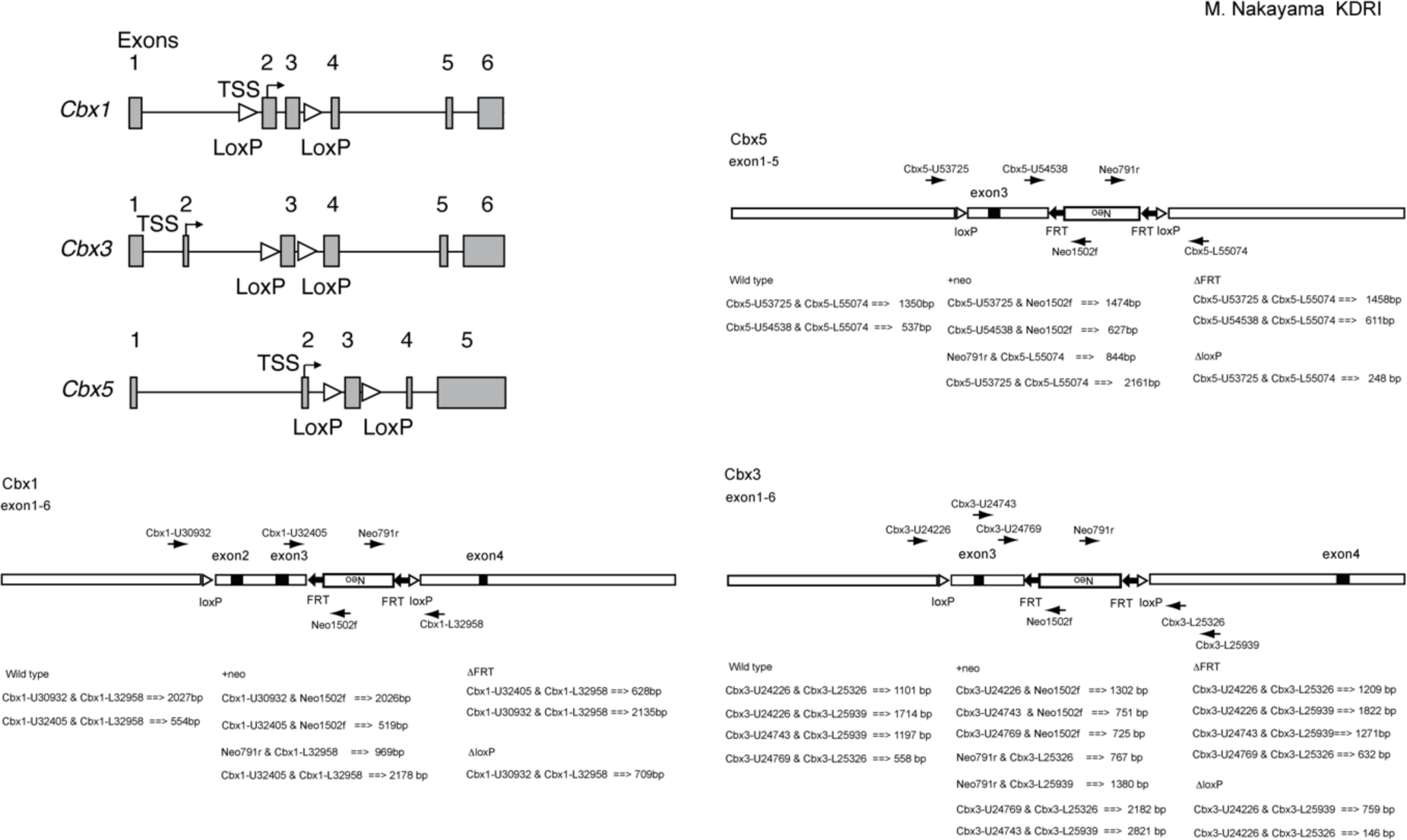
Targeting of *Cbx1*, *Cbx3*, & *Cbx5* alleles in HP1cTKO ESCs

## References

1. Madabhushi, R. et al. Activity-Induced DNA Breaks Govern the Expression of Neuronal Early-Response Genes. Cell 161, 1592–1605 (2015).

2. Wu, W. et al. Neuronal enhancers are hotspots for DNA single-strand break repair. Nature 593, 440–444 (2021).

3. Stefanatos, R. & Sanz, A. The role of mitochondrial ROS in the aging brain. FEBS letters 592, 743–758 (2018).

4. Lu, T. et al. Gene regulation and DNA damage in the ageing human brain. Nature 429, 883–891 (2004).

5. Koen, J. D. & Rugg, M. D. Neural Dedifferentiation in the Aging Brain. Trends in Cognitive Sciences 23, 547–559 (2019).

6. Zampieri, M. et al. Reconfiguration of DNA methylation in aging. Mechanisms of Ageing and Development 151, 60–70 (2015).

7. Horvath, S. DNA methylation age of human tissues and cell types. Genome Biol. 14, R115 (2013).

8. De Cecco, M. et al. Genomes of replicatively senescent cells undergo global epigenetic changes leading to gene silencing and activation of transposable elements. Aging Cell 12, 247–256 (2013).

9. Zhang, W. et al. A Werner syndrome stem cell model unveils heterochromatin alterations as a driver of human aging. Science 348, 1160–1163 (2015).

10. De Cecco, M. et al. Transposable elements become active and mobile in the genomes of aging mammalian somatic tissues. Aging (Albany NY) 5, 867 (2013).

11. Shumaker, D. K. et al. Mutant nuclear lamin A leads to progressive alterations of epigenetic control in premature aging. Proceedings of the National Academy of Sciences 103, 8703–8708 (2006).

12. Ryu, S. H., Kang, K., Yoo, T., Joe, C. O. & Chung, J. H. Transcriptional repression of repeat-derived transcripts correlates with histone hypoacetylation at repetitive DNA elements in aged mice brain. Experimental Gerontology 46, 811–818 (2011).

13. Nellåker, C. et al. Transactivation of elements in the human endogenous retrovirus W family by viral infection. Retrovirology 3, 1 (2006).

14. Li, F. et al. Transcriptional derepression of the ERVWE1 locus following influenza A virus infection. Journal of virology 88, 4328–4337 (2014).

15. Lemaître, C., Tsang, J., Bireau, C., Heidmann, T. & Dewannieux, M. A human endogenous retrovirus-derived gene that can contribute to oncogenesis by activating the ERK pathway and inducing migration and invasion. PLOS Pathogens 13, e1006451 (2017).

16. Prudencio, M. et al. Repetitive element transcripts are elevated in the brain of C9orf72 ALS/FTLD patients. Hum. Mol. Genet. 26, 3421–3431 (2017).

17. Dembny, P., et al. Human endogenous retrovirus HERV-K(HML-2) RNA causes neurodegeneration through Toll-like receptors. JCI Insight 5, (2020).

18. van Horssen, J., van der Pol, S., Nijland, P., Amor, S. & Perron, H. Human endogenous retrovirus W in brain lesions: Rationale for targeted therapy in multiple sclerosis. Multiple Sclerosis and Related Disorders 8, 11–18 (2016).

19. Li, W. et al. Human endogenous retrovirus-K contributes to motor neuron disease. Science translational medicine 7, 307ra153–307ra153 (2015).

20. Douville, R., Liu, J., Rothstein, J. & Nath, A. Identification of active loci of a human endogenous retrovirus in neurons of patients with amyotrophic lateral sclerosis. Annals of neurology 69, 141–151 (2011).

21. Frost, B., Hemberg, M., Lewis, J. & Feany, M. B. Tau promotes neurodegeneration through global chromatin relaxation. Nature Neuroscience 17, 357–366 (2014).

22. Chang, Y.-H. & Dubnau, J. Endogenous retroviruses and TDP-43 proteinopathy form a sustaining feedback driving intercellular spread of Drosophila neurodegeneration. Nat Commun 14, 966 (2023).

23. Desplats, P. et al. α-Synuclein Sequesters Dnmt1 from the Nucleus: A NOVEL MECHANISM FOR EPIGENETIC ALTERATIONS IN LEWY BODY DISEASES. Journal of Biological Chemistry 286, 9031–9037 (2011).

24. Chen, V. et al. The mechanistic role of alpha-synuclein in the nucleus: Impaired nuclear function caused by familial Parkinson’s disease SNCA mutations. Human Molecular Genetics 29, 3107–3121 (2020).

25. Geis, F. K. & Goff, S. P. Silencing and Transcriptional Regulation of Endogenous Retroviruses: An Overview. Viruses 12, 884 (2020).

26. Fukuda, K. & Shinkai, Y. SETDB1-Mediated Silencing of Retroelements. Viruses 12, 596 (2020).

27. Bannister, A. J. et al. Selective recognition of methylated lysine 9 on histone H3 by the HP1 chromo domain. Nature 410, 120–124 (2001).

28. Nielsen, P. R. et al. Structure of the HP1 chromodomain bound to histone H3 methylated at lysine 9. Nature 416, 103 (2002).

29. Singh, P. B. & Newman, A. G. On the relations of phase separation and Hi-C maps to epigenetics. R. Soc. open sci. 7, 191976 (2020).

30. Stephan, A. H. et al. A Dramatic Increase of C1q Protein in the CNS during Normal Aging. J Neurosci 33, 13460–13474 (2013).

31. Reichwald, J., Danner, S., Wiederhold, K.-H. & Staufenbiel, M. Expression of complement system components during aging and amyloid deposition in APP transgenic mice. Journal of Neuroinflammation 6, 35 (2009).

32. Aucott, R. et al. HP1-β is required for development of the cerebral neocortex and neuromuscular junctions. The Journal of cell biology 183, 597–606 (2008).

33. Maksakova, I. et al. Distinct roles of KAP1, HP1 and G9a/GLP in silencing of the two-cell-specific retrotransposon MERVL in mouse ES cells. Epigenetics & Chromatin 6, 1–16 (2013).

34. Schotta, G. A silencing pathway to induce H3-K9 and H4-K20 trimethylation at constitutive heterochromatin. Genes & Development 18, 1251–1262 (2004).

35. Takada, Y. et al. HP1γ links histone methylation marks to meiotic synapsis in mice. Development 138, 4207–4217 (2011).

36. Kiefer, L. et al. WAPL functions as a rheostat of Protocadherin isoform diversity that controls neural wiring. Science 380, eadf8440 (2023).

37. Jin, J. et al. CTCF barrier breaking by ZFP661 promotes protocadherin diversity in mammalian brains. 2023.05.08.539838 Preprint at 10.1101/2023.05.08.539838 (2023).

38. Lv, X. et al. Patterned cPCDH expression regulates the fine organization of the neocortex. Nature 612, 503–511 (2022).

39. Souza, P. P. et al. The histone methyltransferase SUV420H2 and Heterochromatin Proteins HP1 interact but show different dynamic behaviours. BMC Cell Biology 10, 41 (2009).

40. Jiang, Y. et al. The methyltransferase SETDB1 regulates a large neuron-specific topological chromatin domain. Nature Genetics 49, 1239–1250 (2017).

41. Zhao, Y.-T. et al. Long genes linked to autism spectrum disorders harbor broad enhancer-like chromatin domains. Genome Res. 28, 933–942 (2018).

42. Riso, V. et al. ZFP57 maintains the parent-of-origin-specific expression of the imprinted genes and differentially affects non-imprinted targets in mouse embryonic stem cells. Nucleic Acids Research 44, 8165–8178 (2016).

43. Wu, X. & Zhang, Y. TET-mediated active DNA demethylation: mechanism, function and beyond. Nat Rev Genet 18, 517–534 (2017).

44. Stevens, B. et al. The Classical Complement Cascade Mediates CNS Synapse Elimination. Cell 131, 1164–1178 (2007).

45. Wu, T. et al. Complement C3 Is Activated in Human AD Brain and Is Required for Neurodegeneration in Mouse Models of Amyloidosis and Tauopathy. Cell Reports 28, 2111–2123.e6 (2019).

46. Hahn, O. et al. Atlas of the aging mouse brain reveals white matter as vulnerable foci. Cell 186, 4117–4133.e22 (2023).

47. Jönsson, M. E. et al. Activation of endogenous retroviruses during brain development causes an inflammatory response. The EMBO Journal 40, e106423 (2021).

48. Karikó, K., Buckstein, M., Ni, H. & Weissman, D. Suppression of RNA Recognition by Toll-like Receptors: The Impact of Nucleoside Modification and the Evolutionary Origin of RNA. Immunity 23, 165–175 (2005).

49. Cañadas, I. et al. Tumor innate immunity primed by specific interferon-stimulated endogenous retroviruses. Nat Med 24, 1143–1150 (2018).

50. Smallwood, A., Esteve, P.-O., Pradhan, S. & Carey, M. Functional cooperation between HP1 and DNMT1 mediates gene silencing. Genes & Development 21, 1169–1178 (2007).

51. Noguchi, H. et al. DNA Methyltransferase 1 Is Indispensable for Development of the Hippocampal Dentate Gyrus. The Journal of Neuroscience 36, 6050–6068 (2016).

52. Hutnick, L. K. et al. DNA hypomethylation restricted to the murine forebrain induces cortical degeneration and impairs postnatal neuronal maturation. Human Molecular Genetics 18, 2875–2888 (2009).

53. Martos, S. N. et al. Two approaches reveal a new paradigm of ‘switchable or genetics-influenced allele-specific DNA methylation’ with potential in human disease. Cell Discov 3, 17038 (2017).

54. Barbot, W., Dupressoir, A., Lazar, V. & Heidmann, T. Epigenetic regulation of an IAP retrotransposon in the aging mouse: progressive demethylation and de-silencing of the element by its repetitive induction. Nucleic Acids Res 30, 2365–2373 (2002).

55. Sanchez, D. et al. Aging without Apolipoprotein D: Molecular and cellular modifications in the hippocampus and cortex. Experimental Gerontology 67, 19–47 (2015).

56. Stilling, R. M. et al. De-regulation of gene expression and alternative splicing affects distinct cellular pathways in the aging hippocampus. Front. Cell. Neurosci. 8, (2014).

57. de Magalhães, J. P., Curado, J. & Church, G. M. Meta-analysis of age-related gene expression profiles identifies common signatures of aging. Bioinformatics 25, 875–881 (2009).

58. Hong, S. et al. Complement and microglia mediate early synapse loss in Alzheimer mouse models. Science 352, 712–716 (2016).

59. Liddelow, S. A. et al. Neurotoxic reactive astrocytes are induced by activated microglia. Nature 541, 481–487 (2017).

60. Perez-Nievas, B. G. & Serrano-Pozo, A. Deciphering the Astrocyte Reaction in Alzheimer’s Disease. Front. Aging Neurosci. 10, (2018).

61. Pastuzyn, E. D. et al. The Neuronal Gene Arc Encodes a Repurposed Retrotransposon Gag Protein that Mediates Intercellular RNA Transfer. Cell 172, 275–288.e18 (2018).

62. Holm, M. M., Kaiser, J. & Schwab, M. E. Extracellular Vesicles: Multimodal Envoys in Neural Maintenance and Repair. Trends in Neurosciences 41, 360–372 (2018).

63. Cocozza, F. et al. Extracellular vesicles and co-isolated endogenous retroviruses from murine cancer cells differentially affect dendritic cells. The EMBO Journal 42, e113590 (2023).

64. Smith, H. L. et al. Astrocyte Unfolded Protein Response Induces a Specific Reactivity State that Causes Non-Cell-Autonomous Neuronal Degeneration. Neuron 105, 855–866.e5 (2020).

65. Bach, E. A. et al. Ligand-Induced Autoregulation of IFN-γ Receptor β Chain Expression in T Helper Cell Subsets. Science 270, 1215–1218 (1995).

66. Hashioka, S., Klegeris, A., Schwab, C. & McGeer, P. L. Interferon-γ-dependent cytotoxic activation of human astrocytes and astrocytoma cells. Neurobiology of Aging 30, 1924– 1935 (2009).

67. Mitchell, T. J. et al. IFN-gamma up-regulates expression of the complement components C3 and C4 by stabilization of mRNA. J. Immunol. 156, 4429–4434 (1996).

68. Chakrabarty, P. et al. IFN-γ Promotes Complement Expression and Attenuates Amyloid Plaque Deposition in Amyloid β Precursor Protein Transgenic Mice. The Journal of Immunology 184, 5333–5343 (2010).

69. Li, A. et al. IFN-γ promotes τ phosphorylation without affecting mature tangles. The FASEB Journal 29, 4384–4398 (2015).

70. Costa-Mattioli, M. & Walter, P. The integrated stress response: From mechanism to disease. Science 368, (2020).

71. Zhang, J. et al. Neurotoxic microglia promote TDP-43 proteinopathy in progranulin deficiency. Nature 588, 459–465 (2020).

72. Labzin, L. I., Heneka, M. T. & Latz, E. Innate Immunity and Neurodegeneration. Annual Review of Medicine 69, 437–449 (2018).

73. Wyss-Coray, T. Ageing, neurodegeneration and brain rejuvenation. Nature 539, 180–186 (2016).

74. Horie, K. et al. Retrotransposons Influence the Mouse Transcriptome: Implication for the Divergence of Genetic Traits. Genetics 176, 815–827 (2006).

75. Bormuth, I. et al. Neuronal Basic Helix-Loop-Helix Proteins Neurod2/6 Regulate Cortical Commissure Formation before Midline Interactions. Journal of Neuroscience 33, 641–651 (2013).

76. Saito, T. In vivo electroporation in the embryonic mouse central nervous system. Nature Protocols 1, 1552 (2006).

77. Zaqout, S. & Kaindl, A. M. Golgi-Cox Staining Step by Step. Front. Neuroanat. 10, (2016).

78. Sholl, D. A. Dendritic organization in the neurons of the visual and motor cortices of the cat. J Anat 87, 387–406.1 (1953).

79. Schindelin, J. et al. Fiji: an open-source platform for biological-image analysis. Nature Methods 9, 676–682 (2012).

80. Koch, S. et al. Atlas registration for edema-corrected MRI lesion volume in mouse stroke models: Journal of Cerebral Blood Flow & Metabolism (2017) doi:10.1177/0271678X17726635.

81. Sharif, J. et al. Activation of Endogenous Retroviruses in Dnmt1−/− ESCs Involves Disruption of SETDB1-Mediated Repression by NP95 Binding to Hemimethylated DNA. Cell Stem Cell (2016) doi:10.1016/j.stem.2016.03.013.

82. Boyle, P. et al. Gel-free multiplexed reduced representation bisulfite sequencing for large-scale DNA methylation profiling. Genome Biology 13, R92 (2012).

83. Dobin, A. et al. STAR: ultrafast universal RNA-seq aligner. Bioinformatics 29, 15–21 (2013).

84. Jin, Y., Tam, O. H., Paniagua, E. & Hammell, M. TEtranscripts: a package for including transposable elements in differential expression analysis of RNA-seq datasets. Bioinformatics 31, 3593–3599 (2015).

85. Durinck, S., Spellman, P. T., Birney, E. & Huber, W. Mapping Identifiers for the Integration of Genomic Datasets with the R/Bioconductor package biomaRt. Nat Protoc 4, 1184–1191 (2009).

86. Robinson, M. D., McCarthy, D. J. & Smyth, G. K. edgeR: a Bioconductor package for differential expression analysis of digital gene expression data. Bioinformatics 26, 139– 140 (2010).

87. Ramírez, F. et al. deepTools2: a next generation web server for deep-sequencing data analysis. Nucleic Acids Research 44, W160–W165 (2016).

88. Babaian, A. et al. LIONS: analysis suite for detecting and quantifying transposable element initiated transcription from RNA-seq. Bioinformatics 35, 3839–3841 (2019).

89. Rusinova, I. et al. Interferome v2.0: an updated database of annotated interferon-regulated genes. Nucleic Acids Res. 41, D1040–1046 (2013).

90. Merico, D., Isserlin, R., Stueker, O., Emili, A. & Bader, G. D. Enrichment Map: A Network-Based Method for Gene-Set Enrichment Visualization and Interpretation. PLOS ONE 5, e13984 (2010).

91. Kucera, M., Isserlin, R., Arkhangorodsky, A. & Bader, G. D. AutoAnnotate: A Cytoscape app for summarizing networks with semantic annotations. F1000Res 5, 1717 (2016).

92. Akalin, A. et al. methylKit: a comprehensive R package for the analysis of genome-wide DNA methylation profiles. Genome Biol. 13, R87 (2012).

93. Gu, Z., Eils, R. & Schlesner, M. Complex heatmaps reveal patterns and correlations in multidimensional genomic data. Bioinformatics 32, 2847–2849 (2016).

94. Krueger, F. & Andrews, S. R. Bismark: a flexible aligner and methylation caller for Bisulfite-Seq applications. Bioinformatics 27, 1571–1572 (2011).

95. Akalin, A., Franke, V., Vlahoviček, K., Mason, C. E. & Schübeler, D. genomation: a toolkit to summarize, annotate and visualize genomic intervals. Bioinformatics 31, 1127– 1129 (2015).

96. Gu, Z., Gu, L., Eils, R., Schlesner, M. & Brors, B. circlize implements and enhances circular visualization in R. Bioinformatics 30, 2811–2812 (2014).

97. Stubbs, T. M. et al. Multi-tissue DNA methylation age predictor in mouse. Genome Biology 18, 68 (2017).

98. Leung, D. et al. Regulation of DNA methylation turnover at LTR retrotransposons and imprinted loci by the histone methyltransferase Setdb1. PNAS 111, 6690–6695 (2014).

